# Design and analysis of a Proportional-Integral-Derivative controller with biological molecules

**DOI:** 10.1101/303545

**Authors:** Michael Chevalier, Mariana Gómez-Schiavon, Andrew Ng, Hana El-Samad

**Affiliations:** Department of Biochemistry and Biophysics, California Institute for Quantitative Biosciences, University of California San Francisco, United States of America; Department of Bioengineering, University of California Berkeley, United States of America; Chan-Zuckerberg Biohub, United States of America; Co-first author

## Abstract

The ability of cells to regulate their function through feedback control is a fundamental underpinning of life. The capability to engineer *de novo* feedback control with biological molecules is ushering in an era of robust functionality for many applications in biotechnology and medicine. To fulfill their potential, feedback control strategies implemented with biological molecules need to be generalizable, modular and operationally predictable. Proportional-Integral-Derivative (PID) control fulfills this role for technological systems and is a commonly used strategy in engineering. Integral feedback control allows a system to return to an invariant steady-state value after step disturbances, hence enabling its robust operation. Proportional and derivative feedback control used with integral control help sculpt the dynamics of the return to steady-state following perturbation. Recently, a biomolecular implementation of integral control was proposed based on an antithetic motif in which two molecules interact stoichiometrically to annihilate each other’s function. In this work, we report how proportional and derivative implementations can be layered on top of this integral architecture to achieve a biochemical PID control design. We illustrate through computational and analytical treatments that the addition of proportional and derivative control improves performance, and discuss practical biomolecular implementations of these control strategies.

## 1 Introduction

We are entering the era of live cell therapeutics in which the immense capabilities of cells can be harnessed and augmented with rationally designed and engineered functionalities to tackle disease wherever it happens in the human body. For example, CAR T-cells are cells engineered to produce chimeric antigen receptors (or CARs) that recognize and attach to antigens on tumor cells, and have been used to cure subsets of blood cancers. The successful clinical use of these cells was a pivotal moment, but one that was also shadowed by safety concerns. In some recipients of this therapy, these powerful engineered immune cells triggered overwhelming and sometimes life threatening immune responses [1,2]. Moreover, once these cells are released into the patient, no recourse is available to influence them further. Other than inhibitory drug treatment to suppress the therapy at a critical time [3], no efficient modulation of the therapy has been implemented. For this reason, engineering feedback control in therapeutic cells represents a crucial next frontier, allowing these cells to continuously monitor their environment and modulate their function accordingly, and hence enable their robust and safe operation.

Synthetic feedback control has also emerged in metabolic engineering as a promising strategy to circumvent important bottlenecks. For instance, feedback control can be used to limit the production of toxic intermediates in a heterologous pathway, or to balance flux between growth and production of metabolites [4–7]. Specialized strategies resembling proportional feedback control have already been implemented to enhance the production of various molecules in genetically modified cells [8–11]. However, widely applicable feedback control strategies, with quantitatively vetted properties and limitations, are still scarce. Biological feedback control designs that are modular, tunable, and with predictable outcomes are crucial to generalize the use of this strategy in metabolic engineering and other biological fields.

In engineered systems, the formulation for feedback control follows a standardized form in which the output *w*(*t*) of a process Φ to be controlled is measured, and then compared against a desired set-point *r*(*t*), generating an error signal *e*(*t*) = *r*(*t*) – *w*(*t*). This error signal is then processed by the control system, which produces an appropriate corrective input *u*(*t*) to the process Φ (Figure 1A). The control system can be designed to achieve different desired characteristics for the corrective action, for example driving the error signal *e*(*t*) as close to zero as possible. Many decades of research in the field of control and dynamical systems provided general control strategies for a wide variety of technological applications [12]. One in particular, *integral control*, has found ubiquitous use because of its ability to achieve a fundamental control action: to drive *e*(*t*) exactly to 0 after a step disturbance to Φ. *Integral control* therefore allows the steady-state output of a system to be insensitive to step perturbations. The mathematical formulation of integral control is simple and relies on generating a control signal *u*(*t*) that is a weighted integral of the error signal, or 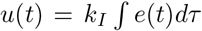. However, integral control is also notorious for its problems with poor damping and instability, which can result in long delays and oscillations before returning to steady-state. This has prompted the widespread use of integral control in combination with other control strategies, such as *proportional* and *derivative* control. *Proportional control* uses the instantaneous error to generate a control signal, that is *u*(*t*) = *k*_*Pe*_(*t*), and by doing so can increase the speed of response. *Derivative control* uses a weighted time derivative of the error signal, 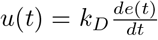, and can increase both the dynamic stability of the controlled system by reducing overshoot and the convergence rate to steady-state. PID-control, a widely used strategy in engineering, capitalizes the benefits of each of these schemes by combining them to generate the control signal, 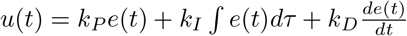 (Figure 1A) [13].

**Fig 1.**
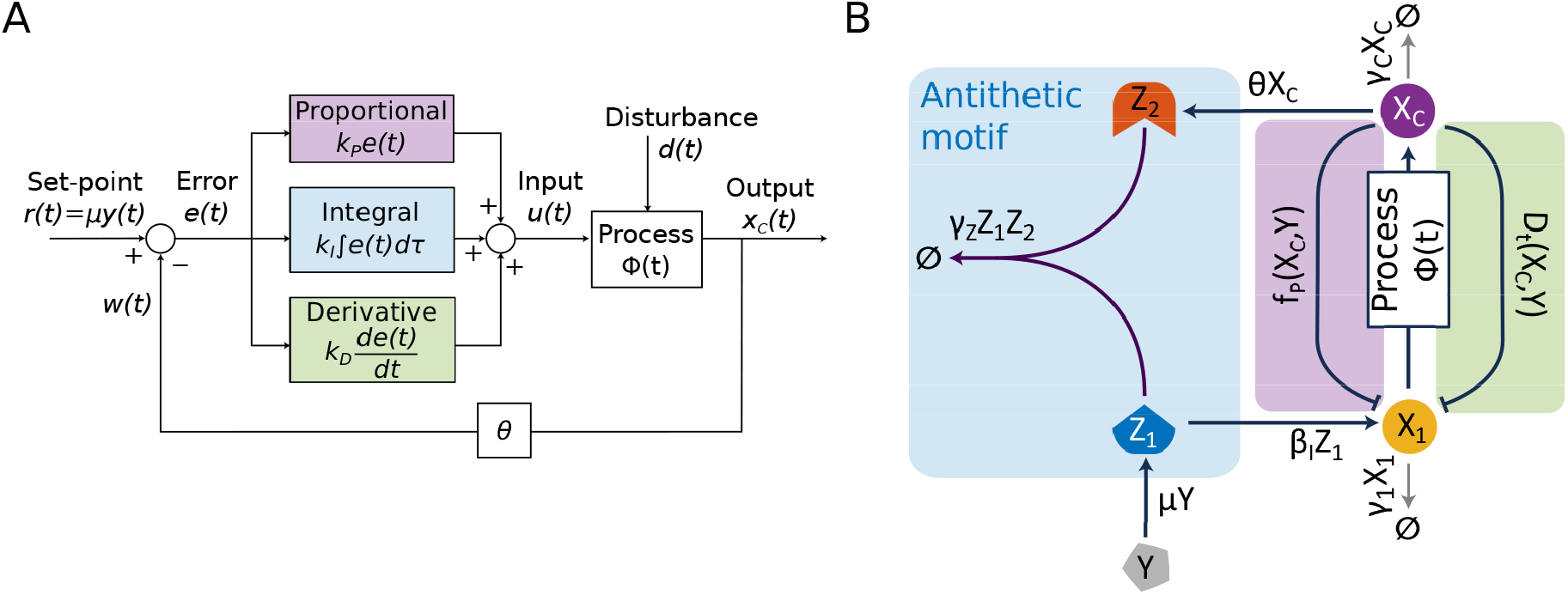
Schematics of an engineering and biomolecular Proportional-Integral-Derivative (PID) controller. (A) Traditional textbook schematic of PID control: Time-dependent regulation error, *e*(*t*) = *r*(*t*) – *w*(*t*) = *μy*(*t*) – *θX*_*C*_(*t*), is continuously computed and processed by the *proportional, integral*, and *derivative* control terms. The control terms are summed up to generate *u*(*t*), the control action that is then fed into the process Φ(*t*). The objective is to eliminate the regulation *error* signal *e*(*t*), driving it to zero after a *disturbance* to the process. (B) Schematic of biomolecular PID control. This design builds on the antithetic motif (blue box) circuit from Briat *et al*. [20]. We add proportional (purple box) and derivative (green box) modules. Like in the traditional PID design, the integral control term (*β*_*I*_*Z*_1_), the proportional control term (*f*_*P*_(*X*_*C*_, *Y*)) and derivative control term (*D*_*t*_(*X*_*C*_, *Y*)) are additive in the actuation of *X*_1_, which is the first molecular species of the controlled process.

Remarkably, many examples of integral feedback have been documented in cellular regulation. Integral feedback was demonstrated to be at work in the *E. coli* chemotaxis circuit, where the percentage of active CheY proteins that are responsible for regulating the bacteria’s tumbling frequency adapts perfectly to step-changes in chemoattractant concentration, maintaining the system’s sensitivity to new concentration changes [14–17]. This was termed *Biochemical Perfect Adaptation*. Other examples of such behavior include the perfect adaptation of the nuclear enrichment of the *S. cerevisiae* MAPK Hog1 after step changes in osmolarity [18], and the control of blood calcium concentration in mammals [19]. In both cases, integral control has been shown to be the structural underpinning of this adaptation.

Because of its widespread successful use in engineering, and its natural occurrence in biology, modular and robust implementations of integral control have been actively pursued in synthetic biology. A breakthrough design of an integral control scheme based on a simple “antithetic motif” was recently reported [20,21]. In this design, two molecular species bind to each other and annihilate each other’s function through this binding (Figure 1B). If one of the “antithetic” molecular species controls the input of a process Φ while the other is dependent on the output of Φ, then it can be mathematically demonstrated that the steady-state value of the output of Φ perfectly adapts regardless of any step perturbation that Φ is subjected to. The antithetic motif used in this configuration therefore implements integral feedback action. An initial experimental proof of concept based on the antithetic feedback motif was recently tested in *E. coli* using *σ* and anti-*σ* factors to implement the antithetic reaction, and the results suggest that this feedback method is indeed able to implement integral control *in vivo* [22]. The next frontier is therefore to design and implement a more general biomolecular PID-control scheme in cells.

In this paper, we present the design of such a biomolecular PID-control system, based on the antithetic integral (I) control motif. Our design of proportional (P) control proceeds through the development of a nonlinear comparator function, while the derivative (D) control design borrows from the biology of the *E. coli* chemotaxis regulatory network [15]. We propose practical biomolecular implementations of these control terms, and illustrate how their addition to integral control improves transient adaptation dynamics. We further show through analytical treatments and numerical simulations how our designs relate to a traditional engineering PID controller, and illustrate how such analogy facilitates the choice of parameters for the P, I, and D control functions as well as their effective weights (*k*_*I*_, *k*_*P*_, *k*_*D*_). Finally, we discuss how the modularity of the proposed PID-control allows easy substitution of different integral controllers, demonstrating that the P and D control terms are robust to the type of integral controller used. Our design paves the way for a modular, general, and robust implementation of PID-control that can greatly benefit applications in metabolic engineering and live cell therapies.

## 2 Results

In a landmark paper by Briat *et al*. [20], a simple integral scheme was proposed to control a general biological process Φ. If the output of the process is *X*_*C*_ and its input (that is actuated by the control signal) is *X*_1_ (Figure 1B), then this scheme is given by:

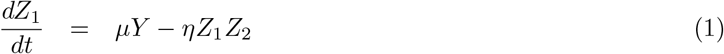

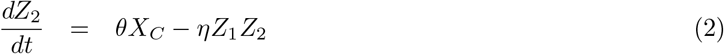

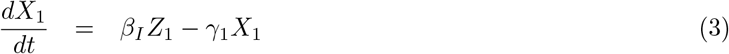

Eqs.(1–2) are produced by an antithetic motif in which two molecules *Z*_1_ and *Z*_2_ bind and annihilate each other’s activity with mass-action kinetics. *Z*_1_ is produced at some rate *μY*, and influences the production of the process-actuated molecule *X*_1_ in Eq.(3). *Z*_2_ is produced at a rate *θΧ*_*C*_, proportional to the output of the process Φ, hence closing the loop (Figure 1B, anthithetic module in blue). The equation describing *Z* = *Z*_1_ – *Z*_2_ is given by:

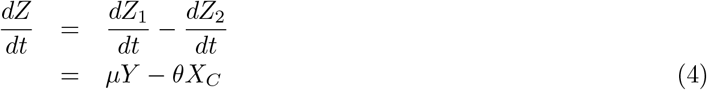

At steady-state, 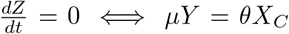. Therefore, the output steady-state is given by 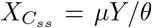, which is obviously independent of the process Φ, and hence is achieved irrespective of Φ and despite any perturbation to its parameters. This is referred to as set-point tracking, since the output at steady-state tracks the input *Y*. Eq.(4) also implies that if the system is at steady-state and a step change in the parameter values of Φ occurs, then *X*_*C*_ will return to its pre-perturbed value after an initial transient. This is the essence of the integral control action. One notes here that *Z* is only a mathematical regulation error quantity and not a physical quantity, as the production of *X*_1_ involves *Z*_1_ and not *Z*_1_ – *Z*_2_. Furthermore, the integral effect is contingent on two general requirements. First, the output *X*_*C*_ must be controllable in a positive manner by the actuated variable *X*_1_ through the process Φ in order to preserve the corrective nature of the feedback through the antithetic motif. For example, saturation should not occur in the observed range of *X*_1_, otherwise *X*_*C*_ cannot be effectively controlled. Second, the disappearance of *Z*_1_ and *Z*_2_ (either through degradation or inactivation) should only be achieved through their mutual annihilation, but not through any other process that affects one but not the other (such as individual degradation of *Z*_1_ and *Z*_2_).

While integral control achieves step disturbance rejection, it is seldom used alone in engineering applications. This is because integral control responds relatively slowly to disturbance, allowing for a large deviation from desired steady-state. This can lead to system instability and oscillations. To illustrate this point, we explore the same simple controlled process Φ described in Briat *et al*. [20] in which *X*_1_ is related to *X*_*C*_ through a gain *β*_*C*_:

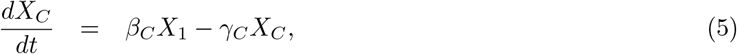

In this system, while perfect disturbance rejection is achieved for some parameter regimes, oscillations also easily emerged with Eqs.(1–3) (which we refer to as I-control) for the controlled process in Eq.(5). This is shown in the left panel of Figure 2A that demonstrates how perfect tracking of the set-point with I-control is only achieved after a long period of damped oscillations. The right panel of Figure 2A shows the I-control system undergoing sustained oscillations following a perturbation in *β*_*C*_, a parameter of the controlled process. Because of this behavior, a commonly used modality in engineering combines integral control with proportional and derivative control, generating Proportional-Integral-Derivative (PID) control. We therefore aim to propose and analyze biochemical implementations of proportional and derivative control that augment and refine the integral control capabilities of the antithetic motif [20]. To achieve this, we propose the following set of equations (Figure 1B):

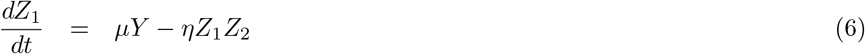

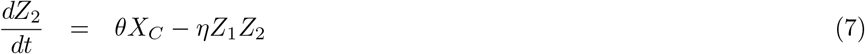

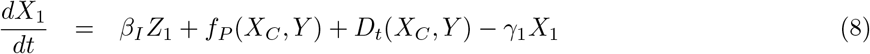

**Fig 2.**
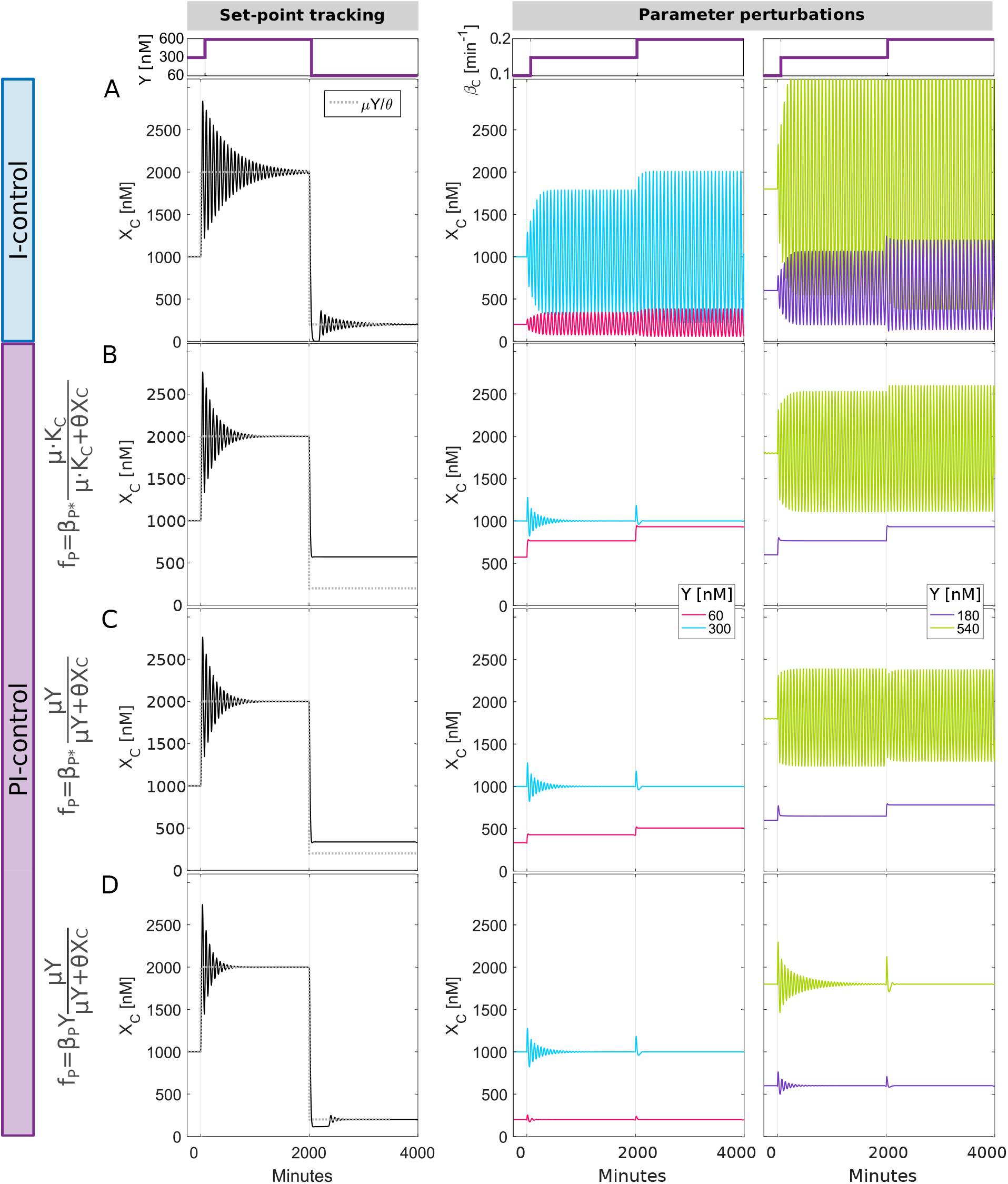
Addition of appropriate proportional control function to integral control improves transient dynamics. (A) Time dynamics of output *X*_*C*_(*t*) following a change in the set-point *Y* (left panel) or in process parameter *β*_*C*_ (right panel). The outcome of *β*_*C*_ perturbation is also shown for different values of *Y* = [60, 180, 300, 540] nM. Simulations are shown for process in Eq.(5) and controller in Eqs.(6–8) of main text. For I-control, *f*_*P*_(*X*_*C*_, *Y*) = 0 and *D*_*t*_(*X*_*C*_, *Y*) = 0 in these equations. (B-D) Time dynamics of output *X*_*C*_(*t*) for different proportional control *f*_*P*_ functions (PI-control, *D*_*t*_(*X*_*C*_, *Y*) = 0). We position *X*_*C*_ at the same value for all control regimes tested for a chosen set-point value of *Y* = 300 nM, hence allowing for a controlled subsequent comparison. See Table 1 for parameter values used in each simulation.

**Table 1.** Simulation parameter values.

| | $\beta_I$ | $\beta_P$ | $\beta_D$ | $\mu$ | $\eta$ | $\theta$ | $\gamma_1$ | $\beta_C$ | $\gamma_C$ | $K_C$ | $\beta_{P*}$ |
| --- | --- | --- | --- | --- | --- | --- | --- | --- | --- | --- | --- |
| **Fig.** | $\text{min}^{-1}$ | $\text{min}^{-1}$ | $\text{min}^{-1}$ | $\text{min}^{-1}$ | $\text{nM}^{-1}$<br>$\text{min}^{-1}$ | $\text{min}^{-1}$ | $\text{min}^{-1}$ | $\text{min}^{-1}$ | $\text{nM}$ | $\text{nM}$ | $\text{nM}$<br>$\text{min}^{-1}$ |
| 2A | 0.06 | 0 | 0 | 1 | 0.01 | 0.3 | 0.1 | $0.1^\ddagger$ | 0.1 | — | — |
| 2B | 0.06 | — | 0 | 1 | 0.01 | 0.3 | 0.1 | $0.1^\ddagger$ | 0.1 | 300 | 90 |
| 2C | 0.06 | — | 0 | 1 | 0.01 | 0.3 | 0.1 | $0.1^\ddagger$ | 0.1 | — | 90 |
| 2D | 0.06 | 0.3 | 0 | 1 | 0.01 | 0.3 | 0.1 | $0.1^\ddagger$ | 0.1 | — | — |
| 3A | 0.06 | 0 | 0.56 | 1 | 0.01 | 0.3 | 0.1 | $0.1^\ddagger$ | 0.1 | — | — |
| 3B | 0.06 | 0 | 0.56 | 1 | 0.01 | 0.3 | 0.1 | $0.1^\ddagger$ | 0.1 | — | — |
| 3C | 0.06 | 0 | 0.56 | 1 | 0.01 | 0.3 | 0.1 | $0.1^\ddagger$ | 0.1 | — | — |
| 3D | 0.06 | 0.15 | 0.28 | 1 | 0.01 | 0.3 | 0.1 | $0.1^\ddagger$ | 0.1 | — | — |
| 4 | 0.06 | 0.15 | 0.28 | 1 | 0.01 | 0.3 | 0.1 | 0.1 | 0.1 | — | — |
| 6A | $0.0576^*$ | 0 | 0 | 1 | 0.01 | 0.3 | 0.1 | $0.1^\ddagger$ | 0.1 | — | — |
| 6B | $0.0576^*$ | 0.3 | 0 | 1 | 0.01 | 0.3 | 0.1 | $0.1^\ddagger$ | 0.1 | — | — |
| 6C | $0.0576^*$ | 0 | 0.6 | 1 | — | 0.3 | 0.1 | $0.1^\ddagger$ | 0.1 | — | — |
| | $\gamma_{A_0}$ | $\beta_A$ | $\gamma_A$ | $\gamma_M$ | $K_A$ | $K_M$ | $\beta_{M*}$ | $\beta_M$ | $\beta_Z$ | $\gamma_Z$ | $K_Z$ |
| **Fig.** | $\text{min}^{-1}$ | $\text{min}^{-1}$ | $\text{min}^{-1}$ | $\text{min}^{-1}$ | $\text{nM}$ | $\text{nM}$ | $\text{nM}$<br>$\text{min}^{-1}$ | $\text{min}^{-1}$ | $\text{min}^{-1}$ | $\text{min}^{-1}$ | $\text{nM}$ |
| 2A | — | — | — | — | — | — | — | — | — | — | — |
| 2B | — | — | — | — | — | — | — | — | — | — | — |
| 2C | — | — | — | — | — | — | — | — | — | — | — |
| 2D | — | — | — | — | — | — | — | — | — | — | — |
| 3A | 0.1 | 1.5 | 1.5 | 1.5 | 1 | 1 | — | 0.4167 | — | — | — |
| 3B | 0.1 | 1.5 | 1.5 | 1.5 | 1 | 1 | 125 | 0.4167 | — | — | — |
| 3C | 0.1 | 1.5 | 1.5 | 1.5 | 1 | 1 | — | 0.4167 | — | — | — |
| 3D | 0.1 | 1.5 | 1.5 | 1.5 | 1 | 1 | — | 0.4167 | — | — | — |
| 4 | 0.1 | 1.5 | 1.5 | 1.5 | 1 | 1 | — | 0.4167 | — | — | — |
| 6A | 0.1 | 1.5 | 1.5 | 1.5 | 1 | 1 | — | 0.4167 | 4 | 2 | 1 |
| 6B | 0.1 | 1.5 | 1.5 | 1.5 | 1 | 1 | — | 0.4167 | 4 | 2 | 1 |
| 6C | 0.1 | 1.5 | 1.5 | 1.5 | 1 | 1 | — | 0.4167 | 4 | 2 | 1 |

$^\ddagger$ Unless directly perturbed (e.g. $\beta_C = [0.1, 0.15, 0.2] \text{min}^{-1}$ ).
$^*$ Corresponds to $\beta_{I*}$ parameter in the alternative PID system (see Section 2.5).

The design upgrades actuation of *X*_1_ with two additional terms, *f*_*P*_(*X*_*C*_, *Y*) for proportional control and *D*_*t*_(*X*_*C*_, *Y*) for derivative control. In the same way that the proportional, integral and derivative control terms are summed up in the engineering PID diagram (Figure 1A), they are summed up in the equation for *X*_1_. We next present designs and analyses that define these terms and their implementation with biomolecules.

### 2.1 Design of a proportional control term

A traditional implementation of proportional control in the engineering sense would require an explicit computation of the tracking error, given by *e*(*t*) = *μΥ* – *θX*_*C*_(*t*) in the context of Figure 1B. Since the outcome of any operation implemented with biological molecules is another molecule, computation of an error signal with biomolecules of this type can only generate a positive quantity. Following the example of the I-control scheme, we will therefore design a proportional control function that acts implicitly on this error, without explicitly computing it.

The first consideration in the design is that for negative feedback (*X*_*C*_ is a negative regulator in the tracking error), the proportional control term *f*_*P*_(*X*_*C*_, *Y*) must be an inhibitory function of *X*_*C*_. One option is to use a traditional Michaelis-Menten negative feedback function, 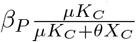, which captures transcriptional repression of *X*_1_ by *X*_*C*_. Here, *β*_*Ρ*_ represents the maximum synthesis rate in the absence of the inhibitor *X*_*C*_, and *μΚ*_*C*_ is a constant related to the affinity or strength of the inhibitor (smaller *K*_*C*_ results in stronger inhibition for the same concentration of *X*_*C*_). We include *μ* and *θ* in the representation of this function in order to keep consistency of notation as it relates to the antithetic feedback description (Figure 1B). This type of transcriptional inhibition function has been explored in natural occurrences of cellular feedback, and previously used in synthetic biology applications [23,24]. However, this function does not explicitly depend on *Y* unlike the tracking error *e*(*t*). This disengagement between the set-point and proportional term limits the dynamic range in a *Proportional-Integral* control system (*D*_*t*_(*X*_*C*_, *Y*) = 0 in Eqs.(6–8)). Specifically, at low reference set-point *Y* values (including *Y* = 0), both perfect tracking behavior and perfect adaptation following a change in *β*_*C*_ are lost (Figure 2B). This is because in this regime, the proportional term introduces basal synthesis of *X*_1_, creating a basal level of *X*_*C*_ and hence of *Z*_2_. At the same time, *μΥ* is too low so that insufficient *Z*_1_ is produced to successfully overcome basal *Z*_2_ level. In fact, *Z*_2_ grows at a positive constant rate since *Z*_1_ is too low to contribute to its annihilation (see Figure S2A). The outcome is that the system is effectively operating in open loop, and the integral control is not satisfactorily active as evidenced by the loss of perfect tracking at low *Y* in Figure 2B. On the other hand, at high *Y* values, excess *X*_*C*_ is produced and the function 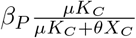 has a small effective value. In this regime, proportional feedback is lost, and the system acts like an I-control scheme (Figure 2B, right panel). As a result, while this function is effective in a range of operation centered around *μΚ*_*C*_, this dynamic range is limited.

A simple upgrade to include dependence both on *Y* and *X*_*C*_ uses the function 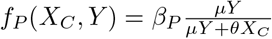. This function has the advantage that it scales with the reference set-point *Y*, allowing for a larger dynamic range before saturation. The result is better tracking at low *Y* and improved damping following perturbations in the parameters of the controlled process such as *β*_*C*_ (Figure 2C). This implementation, however, still incurs a steady-state error from the desired 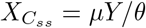 at low *Y* because of the basal production of *X*_*C*_ and *Z*_2_ in this regime, similar to the prior proportional function. Here again, the I-control is not active until *Y* is high enough to engage it. At high values of *Y*, stable oscillations still emerge due to loss of P-control.

We propose the following proportional control strategy that accommodates both low and high *Y* regimes:

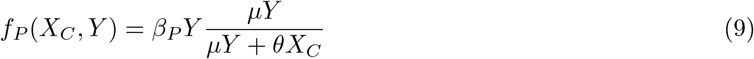

At low *Y* values, *f*_*P*_(*X*_*C*_, *Y*) is proportionally low, reducing the basal synthesis of *X*_*C*_ and *Z*_2_ accordingly and ensuring that I-control is still active. This restores tracking at low *Y* values (Figure 2D, left panel). At high *Y* values, *f*_*P*_(*X*_*C*_, *Y*) also scales accordingly, maintaining the relative contribution of the proportional term to the control system and resulting in improved damping (Figure 2D, right panel). These results were not specific to the deterministic representation that we analyzed, but also held for a stochastic treatment of the system (Figure S1A-B; see Supplemental Information, Section S1, for more details).

The function *f*_*P*_(*X*_*C*_, *Y*) from Eq.(9) can be rearranged to give

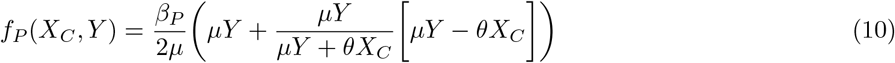

This rearrangement reveals a dependence on the error *e*(*t*) = *μY* – *θX*_*C*_(*t*), which is multiplied by the ratio 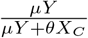 and shifted by *Y* in order to maintain a positive quantity. The superior performance of this functional form of proportional control hinges on this dependence. This is further vetted by analytical arguments presented below in Section 2.3. A possible experimental realization of this function relies on competitive binding to a regulated promoter between *X*_*C*_ and a transcription factor whose activity is proportional to *Y*. This design is discussed in Supplemental Information, Section S2 and Figure Figure S3A.

Finally, we note that although we presented our argument using a function of the form 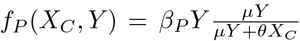 that is positioned at *β*_*P*_*Y*/*2* when the system is at steady-state, our analyses still hold for other less tuned functions. For example, Eq.(9) is derived from a more general biochemical function 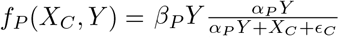 (Supplemental Information, Section S2). When *α*_*P*_ = *μ*/*θ* and *ϵ*_*C*_ = 0, the two functions are the same. If *ϵ*_*C*_ > 0, *f*_*P*_(*X*_*C*_, *Y*) is a weaker proportional feedback function, but this can be compensated by increasing the value of *β*_*P*_ accordingly (Figure S3B). Likewise, if *α*_*P*_ ≫ *μ*/*θ, f*_*P*_(*X*_*C*_, *Y*) is close to *β*_*P*_*Y* at steady state, and the system saturates for negative swings in *X*_*C*_. For *α*_*P*_ ≪ *μ*/*θ, f*_*P*_(*X*_*C*_, *Y*) is a small quantity with very little sensitivity. Here again, these deviations can be compensated for by adjusting the proportional control weight *β*_*P*_ (Figure S3C). However, this benefit of increasing *β*_*P*_ is not unlimited. For a given *β*_*I*_, increasing *β*_*P*_ excessively beyond a certain value may drive *Z*_1_ to be too low to be able to control *Z*_2_, therefore undermining the controller (Figure S2C,F). As with any control design, it is therefore important to design control weights *β*_*I*_ and *β*_*P*_ that avoid this regime given a process and its parameters.

### 2.2 Design of a derivative control term

To design a derivative control term, we drew inspiration from bacterial chemotaxis in which a bacterium’s sensing and adaptation circuit is capable of measuring time derivatives of chemoattractant concentrations as the bacterium swims up or down a gradient [15]. To generate a simple implementation, we adopted a simplified 2-node circuit of the process from Ma *et al*. [25]. Adapting this circuit to our purposes, we propose the following interactions that can perform an approximate time derivative measurement of *X*_*C*_, through the following equations:

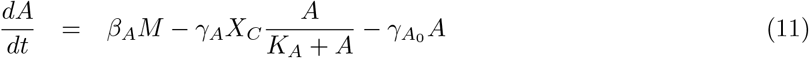

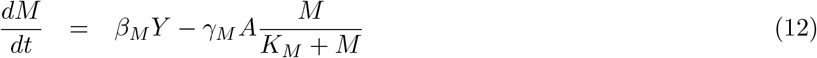

The derivative control term proposed for Eq.(8) is then given by *D*_*t*_(*X*_*C*_, *Y*) = *β*_*D*_*A*, where *A* is the output of the derivative motif in Eqs.(11)–(12). This circuit consists of two molecules, *A* and *M*, where *A* is produced at a rate proportional to *M* and actively degraded by *X*_*C*_. *M* is produced at a rate proportional to the reference signal *Υ* (the signal to be tracked by the controlled process) and actively degraded by *A*. One requirement for the derivative computation through this circuit is that the active degradation terms operate at or near saturation with *K*_*A*_ ≪ *A* and *K*_*M*_ ≪ *M*, over the range of set-point values desired of the process. This results in the following approximate equations:

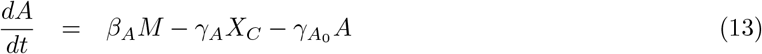

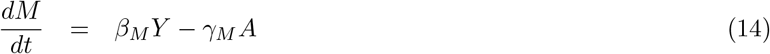

To see how the time-derivative measurement of *X*_*C*_ is achieved, we start by taking the time-derivative of Eq.(13), solving for 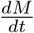 and substituting the resulting expression into Eq.(14), which yields

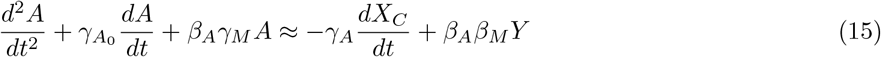

If 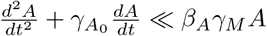, then it follows that

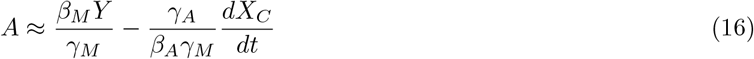

In Eq.(16), *A* is proportional to the negative time derivative of the input *X*_*C*_ plus a steady state value that scales with the reference *Y*. Evidently, the assumptions that lead to Eq.(16) place constraints on the values of 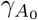, *β*_*Α*_, and *γ*_*Μ*_ given the frequency content of *X*_*C*_, but these are design choices that can be made and tracked (see Supplemental Information, Section S3, and Figure S4 for a procedure and demonstration to fulfill this design and its assumptions). When these design constraints are satisfied, the time dynamics of *A* during its response to either change in reference *Y* or perturbations in *β*_*C*_ generate an accurate measurement of 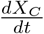 (Figure 3A).

**Fig 3.**
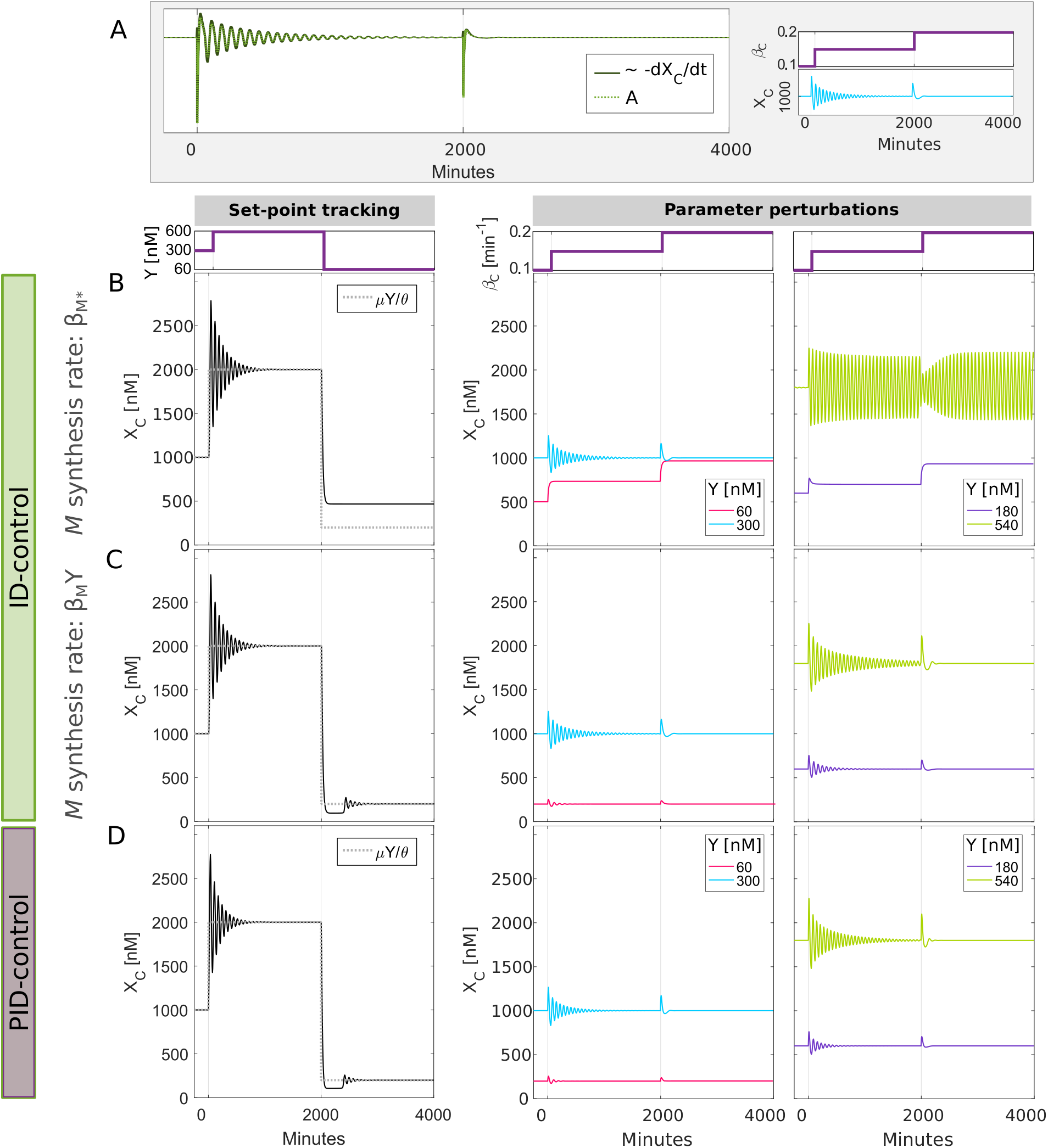
Addition of appropriate derivative control function to integral control improves transient dynamics. (A) Time dynamics of *A* and a numerical approximation of *dX*_*C*_/*dt* for the derivative motif in Eqs.(11–12). Plots shown are for *Y* = 300 nM and perturbations to *β*_*C*_ (see inset). (B) Time dynamics of output *X*_*C*_(*t*) following a change in the set-point *Y* (left panel) or in process parameter *β*_*C*_ (right panel). The outcome of *β*_*C*_ perturbation is also shown for different values of *Y* = [60, 180, 300, 540] nM. Simulations are shown for process in Eq.(5) and controller in Eqs.(6–8) of main text with *f*_*P*_(*X*_*C*_, *Y*) = 0. Derivative function *D*_*t*_(*X*_*C*_, *Y*) is given by *D*_*t*_(*X*_*C*_, *Y*) = *β*_*D*_*A*, where *A* is the output of the derivative motif in Eqs.(11–12), except that in this case, the equation for *M* does not depend on set-point *Y*. (C) Time dynamics of output *X*_*C*_(*t*) for full derivative control design in Eqs.(11–12) with dependence of *M* on set-point *Y*. (D) Simulations for full PID controller under the same conditions as in panels (A-C). See Table 1 for parameter values used in each simulation.

As with proportional control, the addition of a derivative term to the I-control improves its transient response to step *Y* inputs and perturbations in *β*_*C*_ (compare Figure 3C and Figure 2D). Also, like proportional control, the fact that *A* in Eq.(16) scales with *Y* is crucial for the motif to measure the time derivative of *X*_*C*_ over a large dynamic range. To illustrate this point, we compare an Integral-Derivative (ID-, *f*_*P*_(*X*_*C*_, *Y*) = 0) controller in which the production rate of *M* in the derivative calculation motif does not scale with *Y* to the design in which it does (Eq.(12)). The derivative controller that lacks explicit dependence on *Y* also causes loss of perfect tracking (Figure 3B, left panel) and adaptation (Figure 3B, right panel, and Figure Figure S2B) at low *Y* levels. Here again, the I-control is not active until *Y* is high enough to engage it. At high levels of *Y*, oscillations also appear for perturbations in *β*_*C*_. By contrast, basal production at low *Y* and oscillations at high *Y* do not manifest if the synthesis rate of *M* in the derivative control design scales with *Y* (Figure 3C). These results were also robust to biochemical noise (Figure S1C).

Having explored I-, PI- and ID-control, we can now combine all three terms to obtain PID control. Figure 3D shows how the inclusion of both P and I also improves performance in a manner that is similar to that of the PI and ID cases (Figure 2D and Figure 3C, respectively). However, here again, care should be taken in picking the values of the proportional weight *β*_*Ρ*_ and derivative weight *β*_*D*_. If they are too large compared for a given *β*_*I*_, these control terms might undermine the integral control (Figure S2C,D). In Section 2.4, we further explore how the integral and derivative terms affect the performance of the PID controller using a more complicated process to be controlled.

### 2.3 Perturbation analysis of nonlinear PID-control design provides analytical support for the design

To provide an analytical interpretation for the proposed PID controller, we apply linear perturbation theory to Eqs.(6–9,11–12). Even though the simulations above were for the particular controlled process in Eq.(5), the linear analysis is presented for any general controllable process, allowing us to compare our design with a traditional textbook example of a linear PID system extensively used in engineering [26]. In this case, we suppose that the purpose is to control a process whose output is *x*_*C*_(*t*) to a set-point *r*(*t*) (Figure 1A). That is, we want to drive the error *e*(*t*) = *r*(*t*) – *θx*_*C*_(*t*) to zero using a traditional linear PID controller. We will assume that *r*(*t*) = *μy*(*t*) to make this example directly comparable with the biochemical PID controller.

A traditional PID controller uses an input *u*(*t*) into the controlled process that consists of weighted sums of the error (P-control, *k*_*Pe*_(*t*)), the integral of error (I-control, 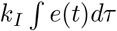), and the time-derivative of error (D-control, 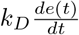) to correct deviations from desired set-point (Figure 1A):

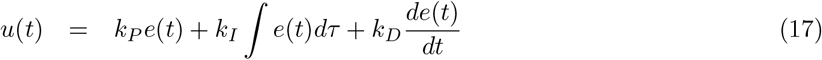

A convenient standard framework for analyzing a linear PID-control system is through frequency domain analysis [26]. A Laplace transform translates time domain signals, e.g. *x*_*C*_(*t*), to frequency domain signals *x*_*C*_(*s*), where *s* is the frequency domain variable. Assuming zero disturbance (*d*(*s*) = 0), *x*_*C*_(*s*) is given by *x*_*C*_(*s*) = *u*(*s*) Φ(*s*), where Φ(*s*) is the process transfer function between the process input *u*(*s*) (the action delivered by the PID controller) and *x*_*C*_(*s*) (Figure 1A). Simple calculations then show that the frequency domain relationship between *r*(*s*) = *μy*(*s*) and *x*_*C*_(*s*) for a traditional PID controller is

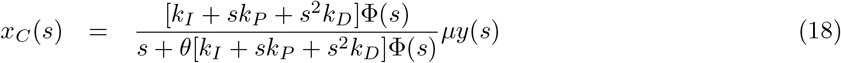

In the frequency domain, steady-state is evaluated at *s* = 0. Assuming that Φ(*s* = 0) ≠ 0, the output at steady-state *x*_*C*_(*s* = 0) is equal to *μy*(*s* = 0)/*θ*, as required for perfect set-point tracking. This is of course only the case when *k*_*I*_ is non-zero, and therefore integral control is necessary.

Evidently, the biochemical controller we are studying is nonlinear. But we can determine its local small-signal properties and relate them to the textbook framework above (Figure 1A and Eq.(18)) using linearization of Eqs.(6–9,11–12) around a steady-state 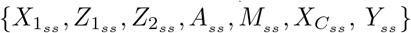 achieved for a desired set-point. This traditional treatment (presented in detail in Supplemental Information, Section S4) generates equations that hold locally for the behavior of the deviation {*x*_1_(*t*), *z*_1_(*t*), *z*_2_(*t*), *a*(*t*), *m*(*t*), *x*_*C*_(*t*)} from the steady-state values as a result of small perturbations to the system. The total solution is the steady state solution plus the perturbed solution; for example, the time-dependent solution for *X*_*C*_(*t*) would be 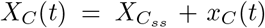. Likewise the input *Y*(*t*) would be *Y*(*t*) = *Y*_*ss*_ + *y*(*t*). Simulations of the linearized controller compared favorably to the nonlinear PID system for small to moderate perturbations (Figure 4 for set-point tracking, and Figure S5A-B for parameter perturbations). Evidently, bigger differences between linearized and nonlinear systems were present for larger perturbations. However these differences were muzzled by adding the quadratic term *ηz*_1_(*t*)*z*_2_(*t*) of the antithetic reaction to the linearized system, suggesting that this is the most impactful nonlinearity in the design for parameter regime used.

**Fig 4.**
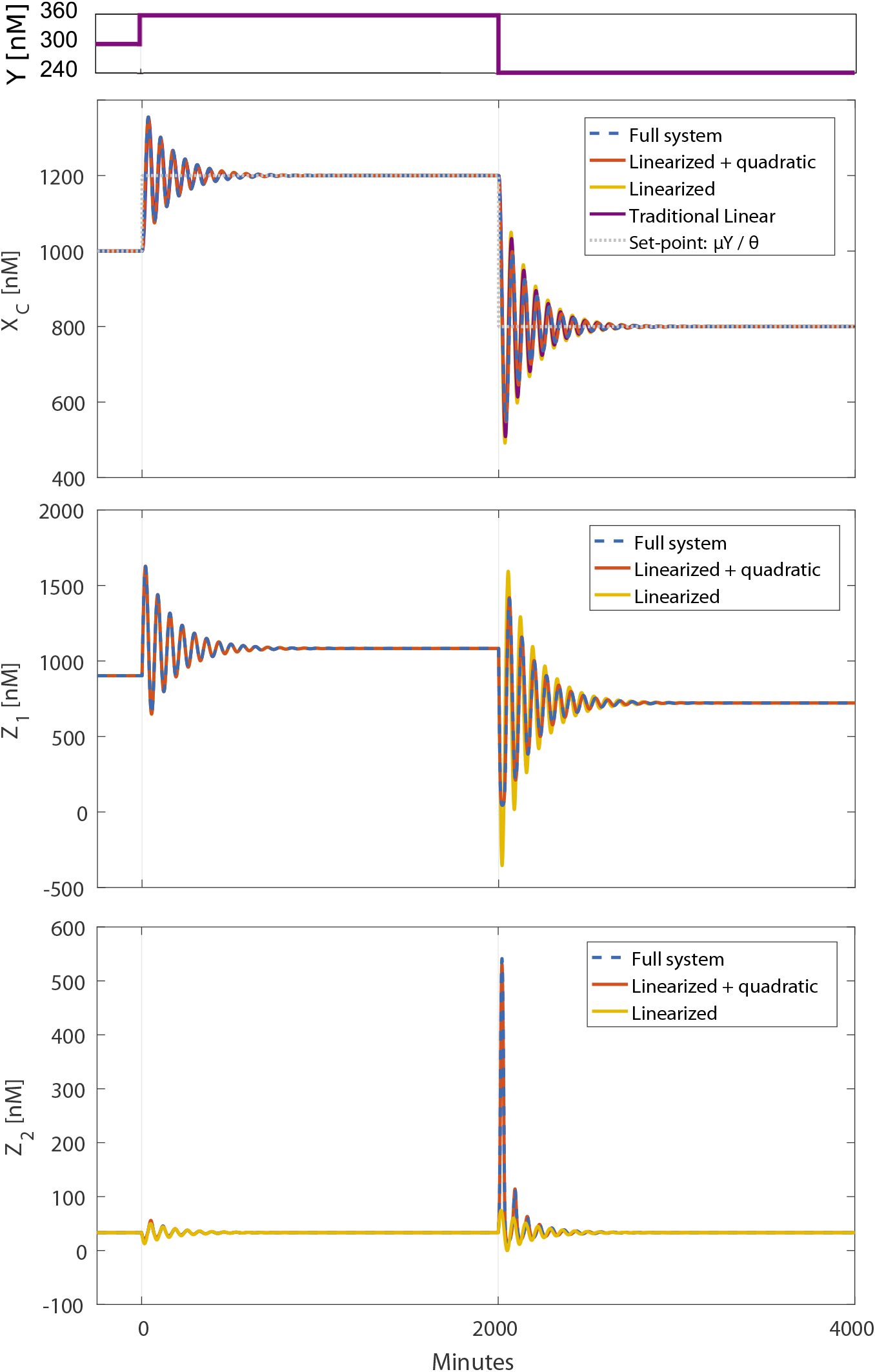
Comparison of nonlinear PID control to its linearized equations in response to perturbations in set-point *Y*. The time dynamics of *X*_*C*_, *Z*_1_ and *Z*_2_ are plotted for the full model (dashed blue, Eqs.(5–8,11–12)), the linearized model (yellow, Eq.(s9)), and the linearized model plus quadratic correction term, *ηz*_1_(*t*)*z*_2_(*t*) (red). Models were simulated with *β*_*I*_ = 0.06 min^−1^, *β*_*Ρ*_ = 0.15 min^−1^ and *β*_*D*_ = 0.28 min^−1^. All other parameters are listed in Table 1. *Y* values are shown in the top panel.

Using the linearized equations for the biomolecular controller, we can investigate how the transfer function between *y*(*s*) and *x*_*C*_(*s*) compares with the traditional PID. The frequency domain transfer function of the linearized biochemical PID controller (see Supplemental Information, Section S4, for derivation) is given by:

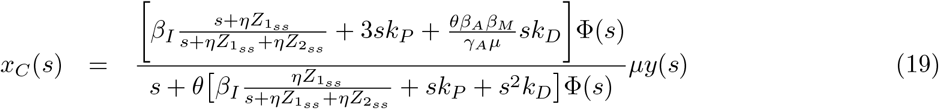

For the process used in this paper (see Eq.(5)), the transfer function between the process input *u*(*s*) and *x*_*C*_(*s*) is 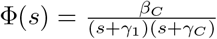. Evidently, at steady state, *x*_*C*_(*s* = 0) = *μy*(*s* = 0)/*θ*, consistent with the full nonlinear system. In this function, *k*_*P*_ is the proportional control gain and *k*_*D*_ is the derivative control gain, which are given by 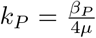 (with 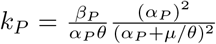, for *α*_*P*_ ≠ *μ*/*θ* in a general proportional term), and 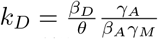, respectively.

While *k*_*D*_ and *k*_*P*_ only depend on parameters and are therefore constant, the integral gain terms in the numerator and denominator of Eq.(19) are a function of the frequency variable *s*. As a result, the linearized antithetic integral control does not exactly map onto a mathematical representation of a traditional linear integral controller. However, these two representations converge under clear constraints on the timescale of the integral controller. Specifically, if *s*_*max*_–the frequency below which resides most of the frequency content of *x*_*C*_(*s*) in the closed loop system-is such that 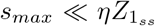, then the function 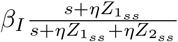 is almost constant over all frequencies below *s*_*max*_. Specifically, when this requirement is met then 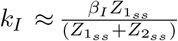. Mechanistically, this could be achieved if binding of the two antithetic molecules that constitute the integral control is much faster than the dynamics of the transcriptional process to be controlled, and this approximation also improves with increasing *Y*, which corresponds to increasing 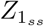 (See supplementary section S4 and Figure S5B). Substituting *k*_*I*_ into the transfer function in Eq.(19) becomes

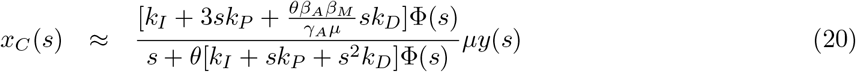

In this approximate transfer function, *k*_*I*_, *k*_*P*_, and *k*_*D*_ are now all constants, and the similarities between this equation and that of the traditional PID controller in Eq.(18) become clear. Eq.(20) has the same denominator (poles of the transfer function) as the traditional PID controller in Eq.(18). But, the two expressions have different numerators (zeros for the transfer function). First, there is a difference in the numerator term that multiplies the proportional gain (*sk*_*P*_ versus 3*sk*_*P*_). The proportional control term in a traditional PID controller acts on the standard tracking error (i.e. *k*_*P*_(*μy* – *θx*_*C*_) in this case). The structure of Eq.(9) generates a linearized proportional control function that acts on a different quantity, manifesting as 3*k*_*P*_*μy* — *k*_*P*_*θx*_*C*_, which is at the root of the difference in the proportional term. This multiplicity of the term *k*_*P*_*y* is in fact the result of *Y* appearing in multiple places in Eq.(9), which leads also to multiplicity in the linearization (see Supplementary Information, Section S4).

Second, the term that multiplies the derivative gain in the biochemical design is given by 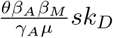, while its counterpart in a traditional design would be *s*^2^*k*_*D*_. This difference can be explained by the fact that the derivative control term in a traditional PID controller acts on the tracking error, i.e. 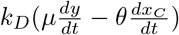, while as we discussed above, the biomolecular implementation of derivative control computes a scaled form of 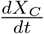 that is also dependent on the reference *Y*, but not its derivative. Even with these differences, and for the designed parameter values we use, the linearized biochemical PID controller showed similar properties as the classical PID controller for set-point tracking (Figure 4) and parameter perturbations (Figure S5A), also see Supplemental Information, Section S4 for simulation implementation details and discussion of an example where some discordance arises.

Eq.(20) and the expressions derived for the effective *k*_*I*_, *k*_*P*_, and *k*_*D*_ provide an analytical framework to discuss the differences in behavior seen for different implementations of the proportional feedback mechanism of Section 2.1. As mentioned above, *k*_*P*_ is constant across set-points for our proposed design of proportional feedback, at least in the linearized regimen. For other proportional functions that were tested and generated narrow dynamic range, that is for 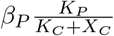 and 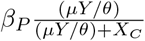, the linearized proportional gains are given by 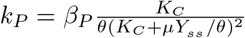 and 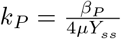, respectively. For both cases, *k*_*P*_ decreases to zero as *Y* increases, indicating that *k*_*P*_ becomes insignificant at higher *Y*, leaving only an I-control like behavior (Figure 2A, right panel). For low *Y*, where these proportional functions caused basal levels of *X*_*C*_ and *Z*_2_, the integral control is not active and *Z*_2_ grows at a positive constant rate since *Z*_1_ is too low to contribute to its annihilation (see Figure S2A). While the other variables in the system reach a steady-state, a transfer function is no longer applicable, nor is *k*_*I*_ defined since it depends on the steady-state value of *Z*_2_. These analytical considerations therefore further support the conclusions reached by numerical analyses of the control system in its nonlinear operation (Figure 2D and Figure 3C).

### 2.4 PID benefits depend on the process to be controlled and PID gains need to be tuned

To move our analysis beyond a simple process that only contains a simple transcriptional step, we consider a more general process connecting *X*_1_ to *X*_*C*_ with delay and negative feedback defined by the following equations

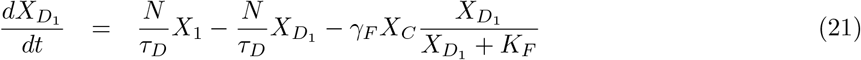

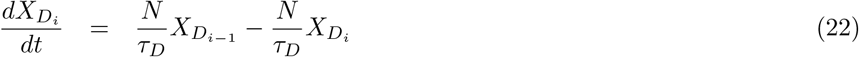

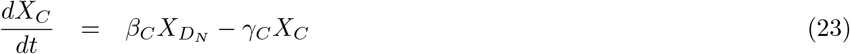

with *X*_1_ still given by Eq.(8). For 2 ≤ *i* ≤ *N* and *γ*_*F*_ = 0, the process is a pure delay of *τ*_*D*_ broken into *N* steps. We now consider two specific process examples. First, when *γ*_*F*_ = 0 min^−1^, *τ*_*D*_ = 20 min, *N* = 2, we obtain a process in which the open loop response for a step change in *Z*_1_ monotonically increases to its new steady state and does not contain oscillations (Figure 5A). Second, for *τ*_*D*_ = 40 min, *N* = 4, and *γ*_*F*_ = 0.2 min^−1^, we generate a process in which a negative feedback loop from *X*_*C*_ onto 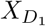 exists. In this case, the open loop process response exhibits damped oscillations (Figure 5B). We explore the benefits of introducing feedback control for the two processes. For the first process, a tuned PI controller can achieve a dynamic response with almost no overshoot (Figure 5A, bottom plot, blue curve). Increasing *k*_*I*_ in this PI controller is detrimental as it adds some overshoot (Figure 5A, bottom plot, red curve) which cannot be corrected by adding a derivative control term (Figure 5A, bottom plot, orange curve). By contrast, for the second process, even a tuned PI controller still generates a closed loop response with a slow oscillating convergence to the set-point (Figure 5B, bottom plot, blue curve). Here, however, addition of a derivative control term improves this transient performance (Figure 5B, bottom plot, orange curve). These results indicate that the benefits of a full PID controller manifest differently for different biological processes and furthermore, that the contributions of the different feedback modalities also need to be tuned and refined based on the specific properties and timescales of the biological process to be controlled. This is similar to considerations that are routinely taken into account for the design and implementation of control strategies in technological systems.

**Fig 5.**
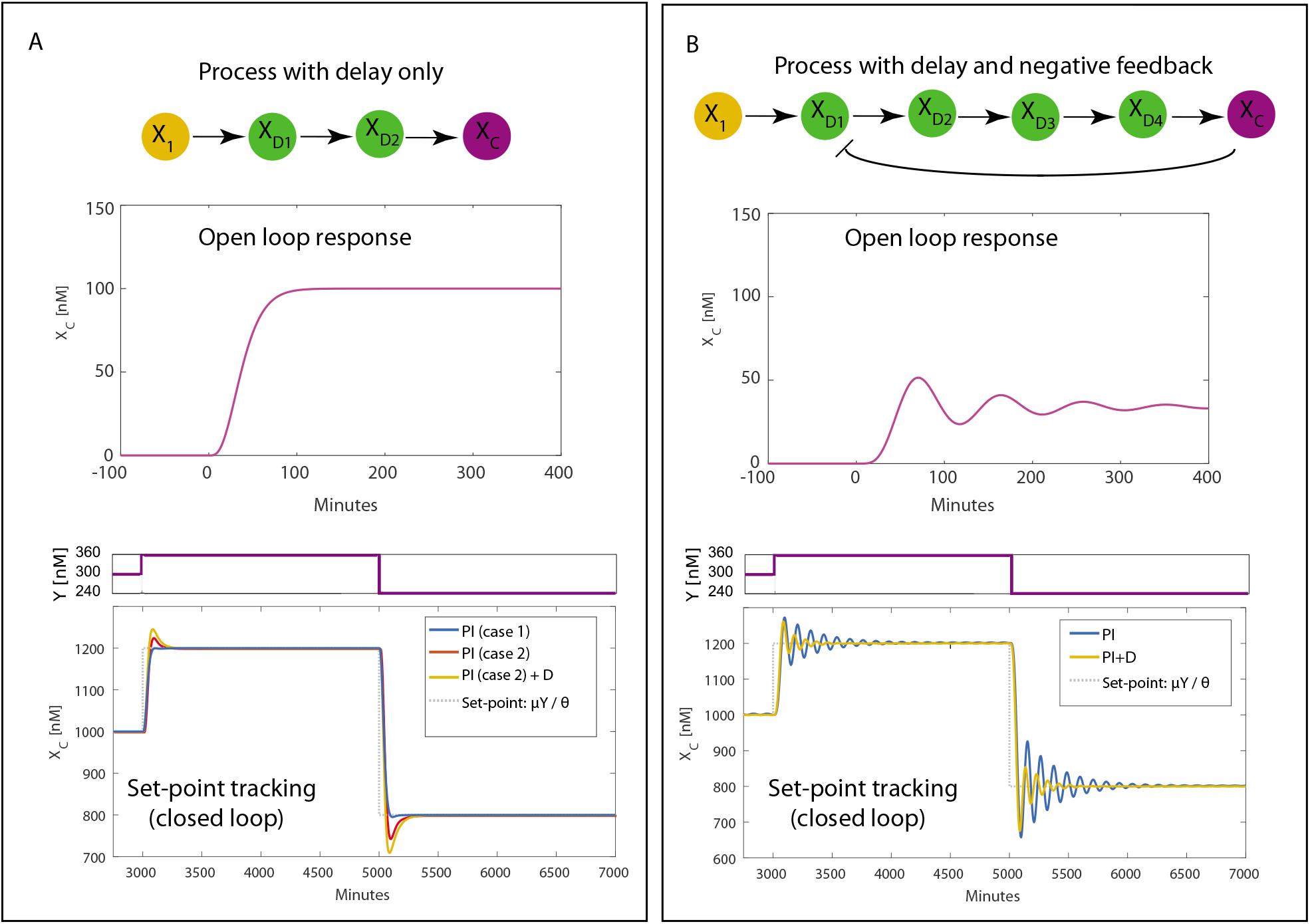
PID benefits depend on the process to be controlled and PID gains need to be tuned accordingly. Panels A-B: Top illustration shows a molecular diagram of the process to be controlled, middle plot shows the open loop response of process to a step change in *Z*_1_, and the bottom plot shows set-point tracking dynamics for different examples of parameters for the feedback controller. (A) Process with just delay (*N* = 2, *τ*_*D*_ = 20 min, and *γ*_*F*_ = 0 min^−1^). Open loop response does not show pronounced oscillations. Tuned PI-controller (case 1: with *k*_*I*_ = 0.00375, *k*_*P*_ = 0.09, and *k*_*D*_ = 0) generates set-point tracking with satisfactory dynamics. Change in *k*_*I*_ (case 2: *k*_*I*_ = 0.0046, *k*_*P*_ = 0.09, and *k*_*D*_ = 0) generates a suboptimal transient response, and addition of derivative control term (case 2 + D: *k*_*I*_ = 0.0046, *k*_*P*_ = 0.09, and *k*_*D*_ = 0.5) does not lead to any improvement. (B) Process with both delay and negative feedback (*N* = 4, *τ*_*D*_ = 40 min, and *γ*_*F*_ = .2 min^−1^). Open loop response shows damped oscillations. Tuned PI controller (*k*_*I*_ = 0.02, *k*_*P*_ = 0.03, and *k*_*D*_ = 0) generates set-point tracking with oscillations. Addition of a derivative control term (PI+D: with *k*_*I*_ = 0.02, *k*_*P*_ = 0.03, and *k*_*D*_ = 4) improves the transient response.

### 2.5 Constructing a PID controller with a different integral controller architecture

We have so far exclusively designed and analyzed proportional and derivative control architectures that are used with the particular antithetic integral control strategy of Briat *et al*. [20]. However, given the additivity of the control terms in Eq.(8), any input from a control design that can implement integral action can be readily used instead. For example, since the proportional control function *f*_*P*_(*X*_*C*_, *Y*) in Eq.(10) has the tracking error encoded within, any biomolecular device that can integrate this function has the potential of implementing integral action, and can therefore be used along with the P and D terms we proposed. To see this, let’s assume that one can construct a variable *Z* that has a rate of change dictated by the following equation:

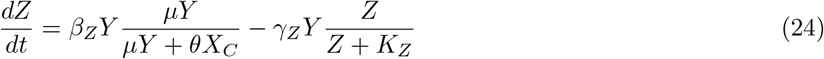

In addition to the function 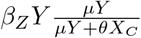 governing the production of *Z*, it is also degraded as a function of *Y*. If this degradation also occurs at saturation (*K*_*Z*_ ≪ *Z*), we have the approximate equation

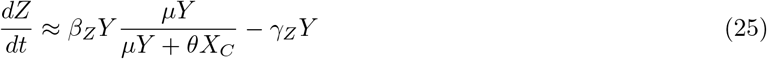

At steady-state,

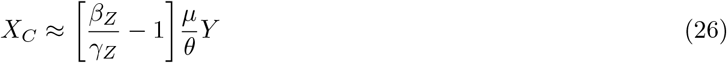

where one sees that *X*_*C*_ is proportional to *Y*, hence implementing perfect tracking. To ensure that both *Y* and *X*_*C*_ are positive, we are evidently constrained by the inequality 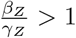.

To implement a PID controller, *Z* can then be input into the control of *X*_1_ in the same fashion as the output of the antithetic motif,

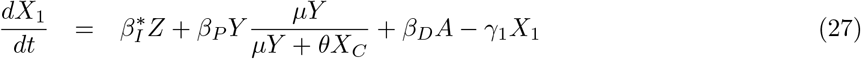

We investigated the use of this new integral controller design as a stand alone, or in PI and ID configurations (replacing Eqs.(6–8) with Eqs.(24,27)). To compare this new design with the antithetic design, we enforced 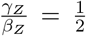, which preserved 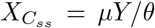. In the linearized regimen, and unlike the antithetic controller, the new design generated a constant integral weight 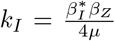 (for derivation of *k*_*I*_ for this new PID see Supplemental Information, Section S4.1), which we set to be equal to the highest generated control weight of the antithetic design. The values for the proportional and derivative controller were kept unchanged and thus both designs share the same *k*_*P*_ and *k*_*D*_. For all perturbations tested, the new integral controller exhibited the same properties as the antithetic implementation (Figure 6A), and its performance was improved by addition of the proportional and derivative control terms (Figure 6B-C, respectively). Taken together, these results indicate that the PID design we propose is modular, and that swapping implementations can be readily done as more designs emerge and are adopted.

**Fig 6.**
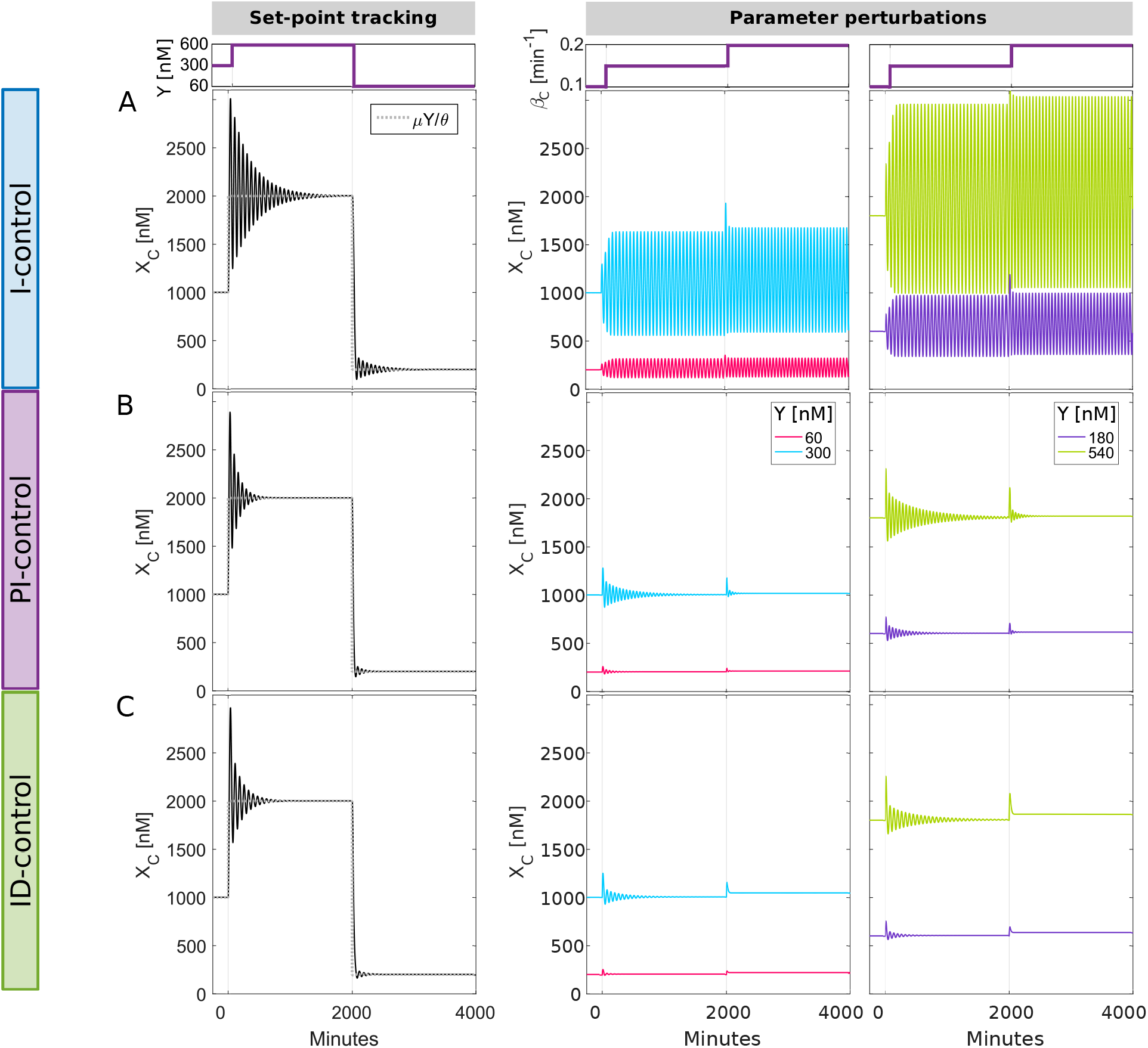
Proportional and derivative control terms improve adaptation dynamics using a new integral controller design. (A) Time dynamics of output *X*_*C*_(*t*) following a change in the set-point *Y* (left panel) or in process parameter *β*_*C*_ (right panel). The outcome of *β*_*C*_ perturbation is also shown for different values of *Y* = [60, 180, 300, 540] nM. Simulations are shown for process in Eq.(5) and new integral controller design in Eqs.(24,27) (I-control; *f*_*P*_(*X*_*C*_, *Y*) = 0 and *D*_*t*_(*X*_*C*_, *Y*) = 0). (B) Time dynamics of output *X*_*C*_(*t*) with *proportional* control (PI-control; *D*_*t*_(*X*_*C*_, *Y*) = 0) using new integral controller design. (C) Time dynamics of output *X*_*C*_(*t*) with *derivative* control (ID-control; *f*_*P*_(*X*_*C*_, *Y*) = 0) using new integral controller design. Parameters used in each panel are listed in Table 1.

## 3 Discussion

Progress in synthetic biology is ushering a future in which predictable interventions in cellular functions and behaviors are becoming possible. Achieving precise control over the intensity, timing, and context of these interventions will generate quantum leaps in many fields, including therapeutic and biotechnological applications. An important cornerstone of such progress is the establishment of capabilities that allow engineered cells, designed for example to deliver a therapeutic function, to regulate this function, deploying it with precision and up-regulating or down-regulating its activity based on the operating environment or internal cellular states. Deploying feedback control of this type will require theoretical and computational studies that deliver plausible design strategies, study their properties and chart their limitations. Importantly, these studies need to take into account the specific properties of the biological substrate, as well as its physical and chemical biological constraints. In doing so, they can transplant crucial notions from the mature field of feedback control field of technological systems into the design and engineering of biology, also updating them as necessary.

Inspired by this notion, we present in this work a design schema for a biology-specific proportional-integral-derivative (PID) controller. PID control has been one of the main workhorses of modern engineering, delivering facile, modular, and tunable control for many applications that we encounter in our every day life, for example our house thermostat. A biochemical PID control strategy endowed with the same properties might also prove to be a general enabling technology for many synthetic biology applications. Our launching design was that for an integral (I) control strategy based on a simple antithetic relationship between two molecules developed by Briat *et al*. [20], which we updated with newly designed biochemical proportional (P) and derivative (D) controllers. Much like their technological counterparts, these additional control terms alleviate the stability constraints of the use of integral controller alone, and provide a malleable and tunable platform to modulate transient dynamics of a controlled system, for example damping down or shortening oscillations. Importantly, through analytical methods, we could relate these designs directly to a traditional formulation of a PID controller, an analogy that facilitates the design and analysis of the biochemical controller based on established theories and practices in other fields. An important feature of the design we propose is its modularity, which we illustrate by exploring PI and ID designs as stand-alone possibilities, and also by swapping the antithetic integral controller with a new implementation inspired by the proportional design strategy (Section 2.5). These properties might prove to be essential for applications in metabolic engineering and cellular therapeutics where different considerations and tradeoffs might be at play, and hence different combinatorial variations of the three terms (P, I, or D) might be needed and appropriate.

While the work we report here presents a design for a general biochemical PID controller, as well as plausible suggestions in terms of molecular building blocks, much work remains to be done to accelerate their implementation and testing. For example, design and implementation of a PID controller for technological systems usually proceeds by experimenting on the system to be controlled in order to determine its properties and hence the controller parameters that might be the most suitable. In our case, success of the controller design relies on identifying the slower timescale of the process and positioning clearly defined parameters of the proportional and derivative controller accordingly. Often time, controller parameters are also fine-tuned in real time during system operation. Carrying out the same process for a biological controller is a formidable challenge, given the long timescales required to build and test in a cell a large number of control strategies or parameter variants. We are hopeful that progress in cellular engineering, as well as more studies that tackle efficient system identification in biological systems will make this cycle more productive, proceeding on timescales that are compatible with rapid deployment of these technologies to various applications.

## Acknowledgements

This work was supported by the Defense Advanced Research Projects Agency, Contract No. HR0011-16-2-0045 to H.E-.S. The content and information does not necessarily reflect the position or the policy of the government, and no official endorsement should be inferred. H.E-.S. is a Chan-Zuckerberg investigator. We would like to thank the members of the El-Samad lab for fruitful discussions.

## Author Contributions

Conceptualization and methodology: MC MGS AN HE. Designed numerical experiments: MC MGS AN. Performed the numerical experiments: MC MGS. Analyzed the data: MC MGS AN HE. Wrote the paper: MC MGS AN HE. Supervision and funding: HE.

## S1 Full chemical equations for PID circuit

The full set of chemical equations for the circuit in Figure 1B, which depict a full PID controller, are

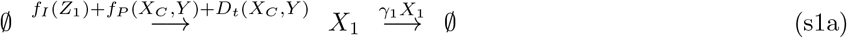

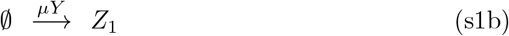

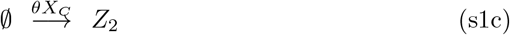

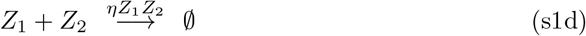

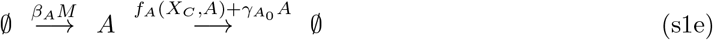

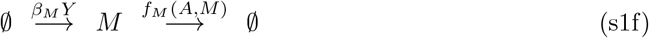

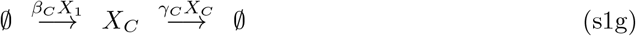

where 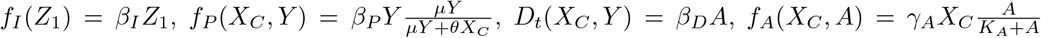, and 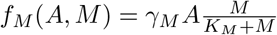.

## S2 Experimental realization of proportional control function

Here, we describe a design for realizing the proportional function *f*_*P*_(*X*_*C*_, *Y*) from Eq.(9) in main text. Many designs can achieve this function, but we focus on one that relies on technologies available in our lab.

The main building block in this design is an inducible transcription factor *T*_*Y*_, which is constitutively expressed at high levels. In the absence of an activating ligand, this transcription factor resides outside the nucleus and it is therefore inactive. *T*_*Y*_ is, for example, a synthetic chimeric transcription factor that has an activating ligand binding domain (LBD) [27]. In the presence of the cognate ligand, *T*_*Y*_ is activated and translocates to the nucleus where it activates a target promoter 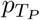. We can modulate the level of active *T*_*Y*_ by adjusting the ligand concentration *Y*, predictably achieving a level of activated transcription factor *Τ*_*Y\**_ equal to *Τ*_*Y\**_ = *G*_*P*_*Y* [27].

We assume that *T*_*Y*\*_ and the output of the process, *X*_*C*_, can compete for a binding site at the promoter 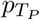 (see Figure S3A). At the promoter, either *T*_*Y*\*_ is bound, *X*_*C*_ is bound, or neither is bound. Therefore, there are three possible promoter states, and transcription can occur only in the active *T*_*Y*_-bound state (*T*_*Y*\*_). Assuming these binding events occur faster than transcription itself, the transcription rate of gene *T*_*P*_ can be computed to be:

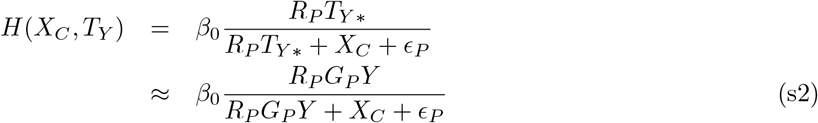

*R*_*P*_ is the ratio of the dissociation constants of *T*_*Y*_ and *X*_*C*_, and *ϵ*_*P*_ is the dissociation constant of *X*_*C*_. Here we assume that *T*_*Y*_ and *X*_*C*_ have the same binding kinetics to 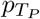. This assumption and the competition at the binding site can be achieved when *T*_*Y*_ and *X*_*C*_ are exactly the same transcription factor protein, but *X*_*C*_ has a crippled transcription activation domain. This can be readily achieved with modular synthetic transcription factors [27]. In this case, *R*_*P*_ ≈ 1. Either way, we set *R*_*P*_*G*_*P*_ = *α*_*P*_ to get

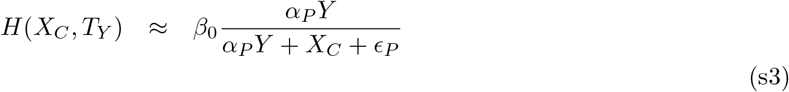

If we choose the minimum set-point to be *α*_*P*_*Y* ≫ *ϵ*_*P*_, then such a design would generate the first proportional function analyzed in the main text that scales with the input *Y*.

To realize the final proportional function, we further assume that the output of the motif above, *T*_*P*_, is a transcription factor whose dynamics are governed by the equation:

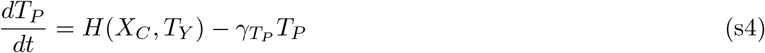

If 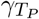 has a fast decay rate, then 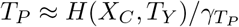. Like *T*_*Y*_, *T*_*P*_ is a cytosolic (inactive) transcription factor that needs to be activated by a ligand to go into the nucleus. Here again, we can modulate externally the active *T*_*P*_ to be proportional to *Y* through changing ligand concentration, setting the activated transcription factor level to 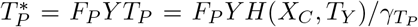. If *X*_1_ is now generated at a rate 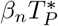 by binding of 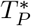 to a cognate promoter, then this generates the proportional control equation:

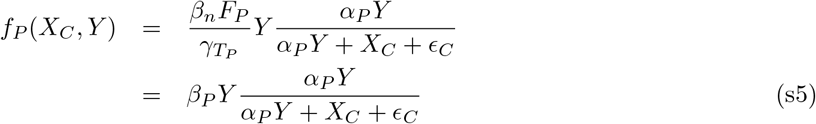

## S3 Realizing the derivative term for the PID-control circuit

Here, we present details of Eq.(16), the approximate time derivative representation of *X*_*C*_ from Section 2.2 in the main text. Transforming Eq.(15) to the frequency domain with frequency variable *s* yields

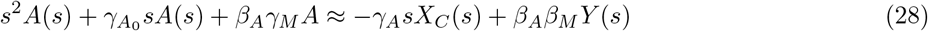

Recall the approximation above (as in Eq.(15)) is from the assumption that the active degradation in the system is occurring at or near saturation, i.e. *K*_*A*_ ≪ *A* and *K*_*M*_ ≪ *M*. Solving for *A*(*s*) yields

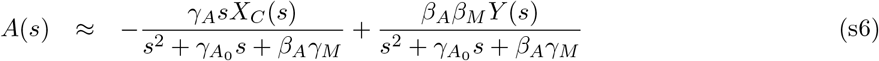

To realize derivative control, we need to design parameter regimes for this motif where *A*(*s*) is approximately proportional to **s*X*_*C*_(*s*), the frequency domain representation of the time derivative of *X*_*C*_.

To do so, we need to design the parameters of the derivative motif so that the poles of the bandpass transfer function 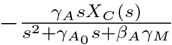 become influential only at high frequencies, that is frequencies that are not relevant to *X*_*C*_(*s*). To insure that, let’s define *F* to be 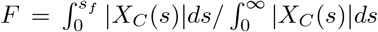. One can evaluate *F* for different values of *s*_*f*_. Let *s*_*max*_ be defined as the frequency at which *F* reaches an acceptable design values, for example *F* = .95. This is the frequency below which resides most of the frequency content of *X*_*C*_(*s*). A conservative design choice is to suppose that the parameter values of the derivative motif are chosen such that 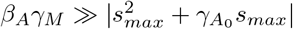, then we obtain

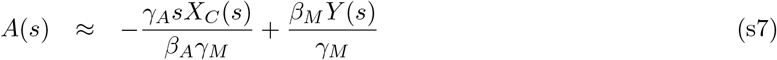

The expression *sX*_*C*_(*s*) in the frequency domain corresponds to 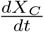 in the time domain. Transforming back to the time-domain yields Eq.(16) in the main text. This result implies that the condition 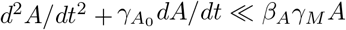 also holds since *d*^2^*A*/*dt*^2^ and 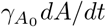 are not present in Eq.(16). It is worth noting here that the design of these parameters should be an iterative process, in which approximate values are picked, the closed-loop assessed, then the parameter values iteratively adjusted in order to correctly position the frequency response of the derivative motif. An illustration of this concept is shown in (Figure S4).

## S4 The control circuit equations linearized about a set-point

Our goal here is to linearize the biochemical PID controller and process to relate it to the textbook linear PID case discussed in Section 2.3. We will derive the frequency-domain transfer function between *μy* and *θx*_*c*_ for our linearized PID biochemical controller.

As a means of evaluating perturbations about a set-point, we linearized Eqs.(6–8) and the simplified derivative equation Eq.(16) around some steady state 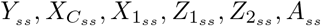. The steady-state values are computed as:

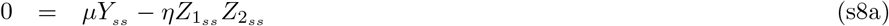

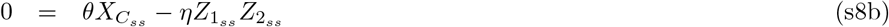

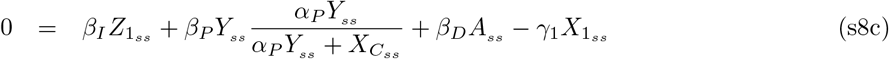

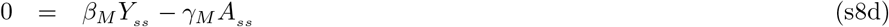

The linearized time-dependent perturbed system is

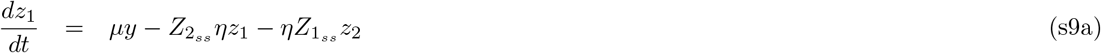

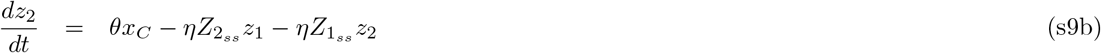

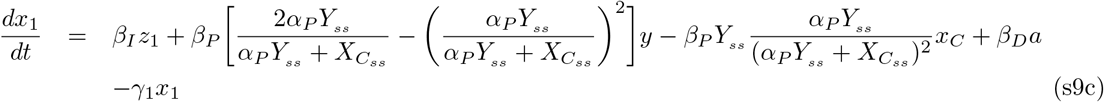

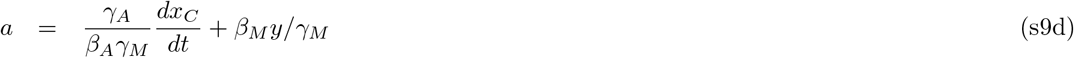

The equations were derived by computing the Jacobian matrix of the nonlinear system and evaluating at steady-state [28]. The approximate solution is locally equal to the steady state solution plus the perturbed solution, for example, the time-dependent solution for *X*_1_ would be 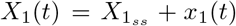. Likewise, the input *Y* would be *Y*(*t*) = *Y*_*ss*_ + *y*(*t*). Transforming this set of linear equations into the frequency domain, we obtain:

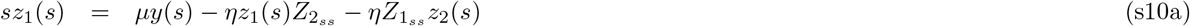

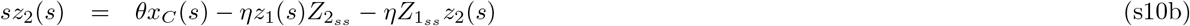

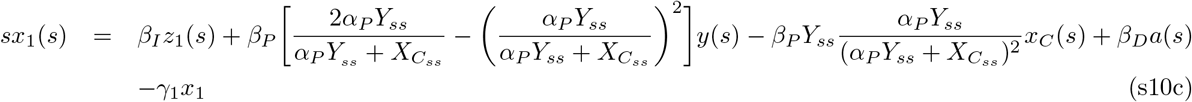

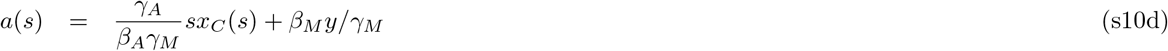

To facilitate notation, we designate 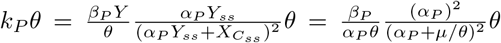. When *α*_*P*_ ≈ *μ*/*θ* to position the proportional controller in its optimal dynamic range, 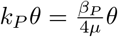. For the rest of the derivation we assume *α*_*P*_ ≈ *μ*/*θ*. We also designate 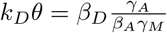 (see Section 2.2). Substituting these constants and the relationship for *a*(*s*) in Eq.(s10d) into Eq.(s10c) the equation becomes

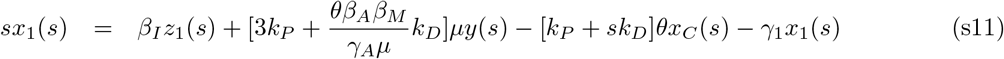

Let *f*(*s*) be the transfer function of the linearized process to be controlled by *x*_1_(*s*). Thus, *x*_*C*_(*s*) = *f*(*s*)*x*_1_(*s*). Using this relationship into Eq.(s11), we obtain after rearrangement

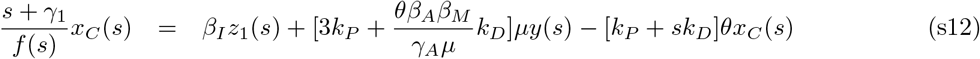

We then use Eqs.(s10a–b) to derive an expression of *z*_1_(*s*) in terms of *x*_*C*_(*s*) and *y*(*s*), and use this expression in equation above to generate:

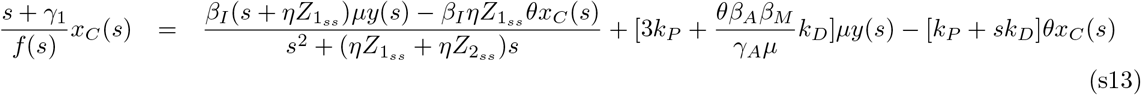

Letting 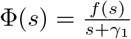, we can rearrange Eq.(s13) to obtain the frequency domain relationship between *x*_*C*_(*s*) and *y*(*s*) of Eq.(19) given by:

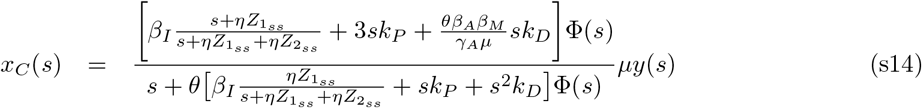

At steady state, i.e. *s* = 0, it is easy to see that *μy*(*s* = 0) = *θx*_*C*_(*s* = 0), consistent with the full nonlinear system. While *k*_*D*_ and *k*_*P*_ are constants that are dependent on parameters, the integral gain terms in the numerator and denominator of Eq.(s14) are a function of the frequency variable *s*. However, if the upper-bound of the frequency content of *x*_*C*_(*s*), i.e. *s*_*max*_ is known, then the parameter *η* of the derivative controller can be designed such that 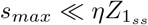. Let’s again define *F* to be 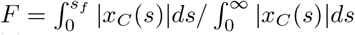 where *F* is the proportion of the area of the integral of the magnitude of *x*_*C*_(*s*) below *s*_*f*_. Let *s*_*max*_ be defined as the frequency at which *F* reaches an acceptable design value, for example *F* = .95. This is the frequency below which resides most of the frequency content of *x*_*C*_(*s*). In the case where 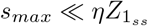, the integral terms in the numerator and denominator can be approximated as 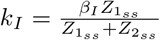. This approximation conforms the numerator and denominator terms that are associated with integral control to the traditional PID expression. Making this approximation in Eq.(19) and letting the integral control weight 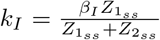 yields Eq.(20) in the main text. For the processes we use in this paper, the highest calculated *s*_*max*_ was approximately 0.25 rad min^−1^. For our lowest input values *Y* = 60 nM, 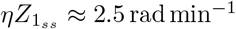, about an order of magnitude larger than *s*_*max*_. Thus, even for this worst case, 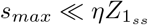. For *Y* = 300 nM, 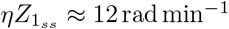. In general for larger *Y*, 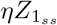 scales approximately with *Y*.

While the analyses above derive linearization and proportional gains for the final form of the proportional control, similar treatment can be extended to the other proportional control functions that are analyzed and compared. For the proportional control function 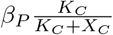, we get 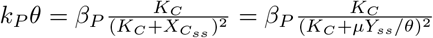. And for the proportional control function 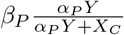, we get 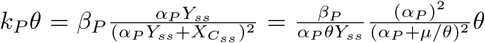 which becomes 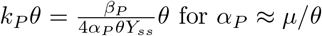.

Finally, to assess the accuracy of the linearization for the parameter values used, we simulated the linearized system and compared it to the full nonlinear system (Figures 4 and Figure S5). In addition, we also simulated a traditional PID controller where we coupled the process equation (Eq.(5) with an *X*_1_ that uses the traditional error terms

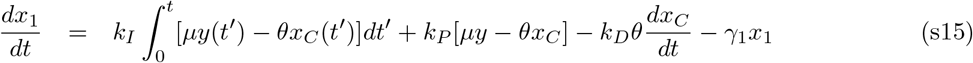

This can be used to compare the linearized biochemical controller to the traditional one, especially scrutinizing the approximation made for *k*_*I*_. Specifically, the transfer function for the traditional PID controller s15 for a step change Δ*β*_*C*_ in *β*_*C*_ is:

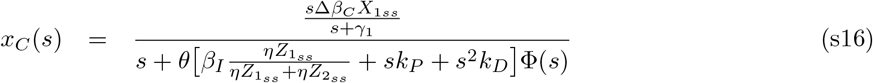

That of the biochemical controller is:

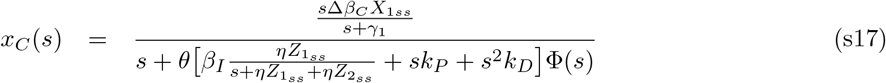

Comparing both, once can see that the integral control gains become more similar to each other as 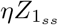 increases, which occurs when *Y* increases. For our examples in this paper, *Y_ss_* = 300 nM is a large enough value (Figure S5C) such that the linearized biochemical PID and the traditional linear PID have a very similar response and hence the linearized biochemical PID is exhibiting a constant *k*_*I*_ in this regime.

For the system simulated to generate Figures 4A and Figure S5A, the traditional linear PID agree well with the linearized biochemical PID. For the case of the process with delay only (Section 2.4), we simulated the linearized version and the traditional linear controller for case 1 from Figure 5A. The full system and the linearized version agree well while the traditional linear controller has a much slower convergence rate (Figure S5D). Given that there is no derivative term in this case, the only source for the differences is the 3*k*_*P*_*y* (linearized version) versus *k*_*P*_*y* (traditional). We simulated the linearized biochemical controller but dividing the 3*k*_*P*_*y* by 3, which resulted in identical results for the traditional and biochemical controller (Figure S5D, linearized (*k*_*P*_*y*)). Interestingly the 3*k*_*P*_*y* term in the biochemical controller helps accelerate convergence relative to the traditional case.

### S4.1 Linearized analysis for PID controller with the new integral implementation

We can carry the same linearization analyses as above to Eq.(25)

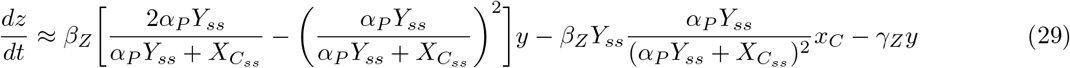

whose frequency domain equation when *α*_*P*_ = *μ*/*θ* and *γ*_*Z*_ = *βz*/2 and 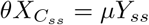 is

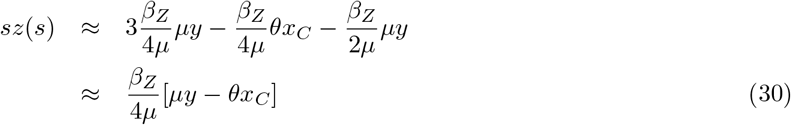

We can then take Eq.(s12) which relates the *x*_*C*_(*s*) to *y*(*s*) for the antithetic PID and replace the antithetic integral control term *β*_1_*z*_1_(*s*) with the new integral control term 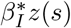 to get

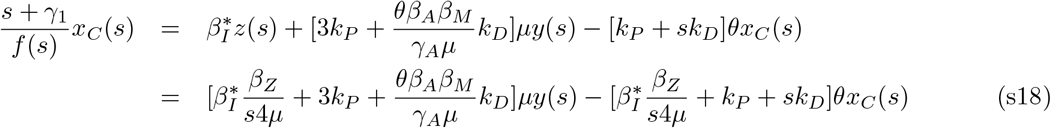

where we set 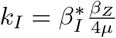 to get the transfer function

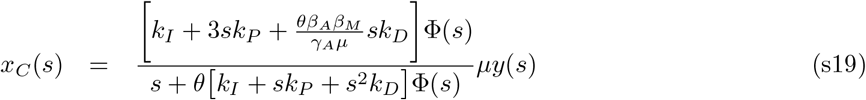

which has exactly the same form as the transfer function for the antithetic system, Eq.(20). The only difference being 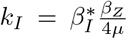 for the new PID while 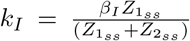 for the antithetic PID, a function of the steady state *Z*_1_ and *Z*_2_ values. When simulating the new PID in Section 2.5, we enforce that 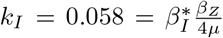, where we set 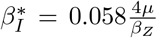 to obtain the same *k*_*I*_ value for both strategies for fair comparison.

**Table S1.** Simulation parameter values in Supplementary Figures.

| | $\beta_I$ | $\beta_P$ | $\beta_D$ | $\mu$ | $\eta$ | $\theta$ | $\gamma_1$ | $\beta_C$ | $\gamma_C$ | $\epsilon_C$ | $\alpha_P$ |
| --- | --- | --- | --- | --- | --- | --- | --- | --- | --- | --- | --- |
| **Fig.** | $\text{min}^{-1}$ | $\text{min}^{-1}$ | $\text{min}^{-1}$ | $\text{min}^{-1}$ | $\text{nM}^{-1}$<br>$\text{min}^{-1}$ | $\text{min}^{-1}$ | $\text{min}^{-1}$ | $\text{min}^{-1}$ | nM | nM<br>$\text{min}^{-1}$ | |
| S1A | 0.06 | 0 | 0 | 1 | 0.01 | 0.3 | 0.1 | $0.1^\ddagger$ | 0.1 | — | — |
| S1B | 0.06 | 0.3 | 0 | 1 | 0.01 | 0.3 | 0.1 | $0.1^\ddagger$ | 0.1 | — | — |
| S1C | 0.06 | 0 | 0.56 | 1 | 0.01 | 0.3 | 0.1 | $0.1^\ddagger$ | 0.1 | — | — |
| S2A | 0.06 | — | 0 | 1 | 0.01 | 0.3 | 0.1 | 0.1 | 0.1 | — | — |
| S2B | 0.06 | 0 | 0.56 | 1 | 0.01 | 0.3 | 0.1 | 0.1 | 0.1 | — | — |
| S2C | 0.06 | 1 | 0 | 1 | 0.01 | 0.3 | 0.1 | 0.1 | 0.1 | — | — |
| S2D | 0.06 | 0 | 1.5 | 1 | 0.01 | 0.3 | 0.1 | 0.1 | 0.1 | — | — |
| S2E | 0.06 | 0.5 | 0.75 | 1 | 0.01 | 0.3 | 0.1 | 0.1 | 0.1 | — | — |
| S2F | 0.06 | 0.3 | 0 | 1 | 0.01 | 0.3 | 0.1 | 0.2 | 0.1 | — | 5 |
| S3B | 0.06 | $0.3^*$ | 0 | 1 | 0.01 | 0.3 | 0.1 | $0.1^\ddagger$ | 0.1 | 500 | — |
| S3C | 0.06 | $0.3^*$ | 0 | 1 | 0.01 | 0.3 | 0.1 | $0.1^\ddagger$ | 0.1 | — | $[\dots]^\dagger$ |
| S5 | 0.06 | 0.3 | 0.28 | 1 | 0.01 | 0.3 | 0.1 | $0.1^\ddagger$ | 0.1 | — | — |
| | $\gamma_{A_0}$ | $\beta_A$ | $\gamma_A$ | $\gamma_M$ | $K_A$ | $K_M$ | $\beta_{M^*}$ | $\beta_M$ | $K_C$ | $\beta_{P^*}$ | |
| **Fig.** | $\text{min}^{-1}$ | $\text{min}^{-1}$ | $\text{min}^{-1}$ | $\text{min}^{-1}$ | nM | nM | nM<br>$\text{min}^{-1}$ | $\text{min}^{-1}$ | nM | nM<br>$\text{min}^{-1}$ | |
| S1A | — | — | — | — | — | — | — | — | — | — |  |
| S1B | — | — | — | — | — | — | — | — | — | — |  |
| S1C | 0.1 | 1.5 | 1.5 | 1.5 | 1 | 1 | — | 0.4167 | — | — |  |
| S2A | — | — | — | — | — | — | — | — | 300 | 90 |  |
| S2B | 0.1 | 1.5 | 1.5 | 1.5 | 1 | 1 | 125 | — | — | — |  |
| S2C | — | — | — | — | — | — | — | — | — | — |  |
| S2D | 0.1 | 1.5 | 1.5 | 1.5 | 1 | 1 | — | 0.4167 | — | — |  |
| S2E | 0.1 | 1.5 | 1.5 | 1.5 | 1 | 1 | — | 0.4167 | — | — |  |
| S2F | — | — | — | — | — | — | — | — | — | — |  |
| S3B | — | — | — | — | — | — | — | — | — | — |  |
| S3C | — | — | — | — | — | — | — | — | — | — |  |
| S5 | 0.1 | 1.5 | 1.5 | 1.5 | 1 | 1 | — | 0.4167 | — | — |  |

$^\ddagger$ Unless directly perturbed (e.g. $\beta_C = [0.1, 0.15, 0.2] \text{ min}^{-1}$ ).
$^*$ Except when stated otherwise.
$^\dagger$ Multiple values: $\alpha_P = [0.5, 0.75, 1, 1.25, 1.5] \frac{\mu}{\theta}$ .
$^\ddagger\ddagger$ Multiple values: $K_A = [1, 10, 100] \text{ nM}$ .
$^\ddagger\ddagger\ddagger$ Multiple values: $K_M = [1, 100, 1000] \text{ nM}$ .

**Figure S1.**
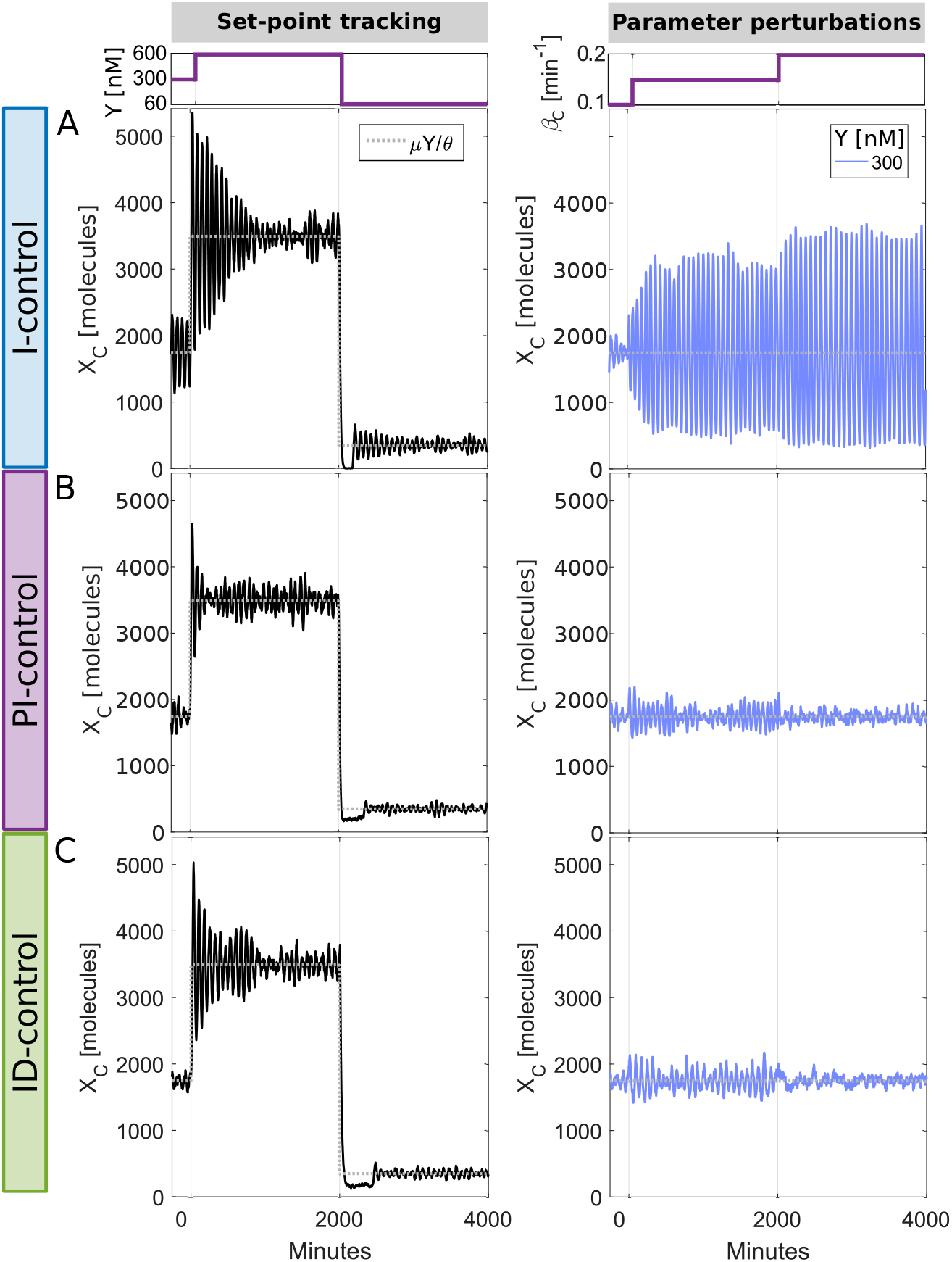
Proportional and derivative control terms improve adaptation dynamics even in the presence of biochemical noise. The model in Eqs.(5–8,11–12) of main text was simulated using the stochastic simulation algorithm. (A) Time dynamics of output *X*_*C*_(*t*) in a single cell following a change in the set-point *Y* (left panel) or in process parameter *β*_*C*_ (right panel) for *Y* = 300 nM when using an integral controller only (I-control; *f*_*P*_(*X*_*C*_, *Y*) = 0 and *D*_*t*_(*X*_*C*_, *Y*) = 0). (B) Same plots as in (A) but after addition of *proportional* control to integral control (PI-control; *D*_*t*_(*X*_*C*_, *Y*) = 0). (C) Same plots as in (A) but after addition of *derivative* control to integral control (ID-control; *f*_*P*_(*X*_*C*_, *Y*) = 0). See Table S1 for parameter values used in each simulation.

**Figure S2.**
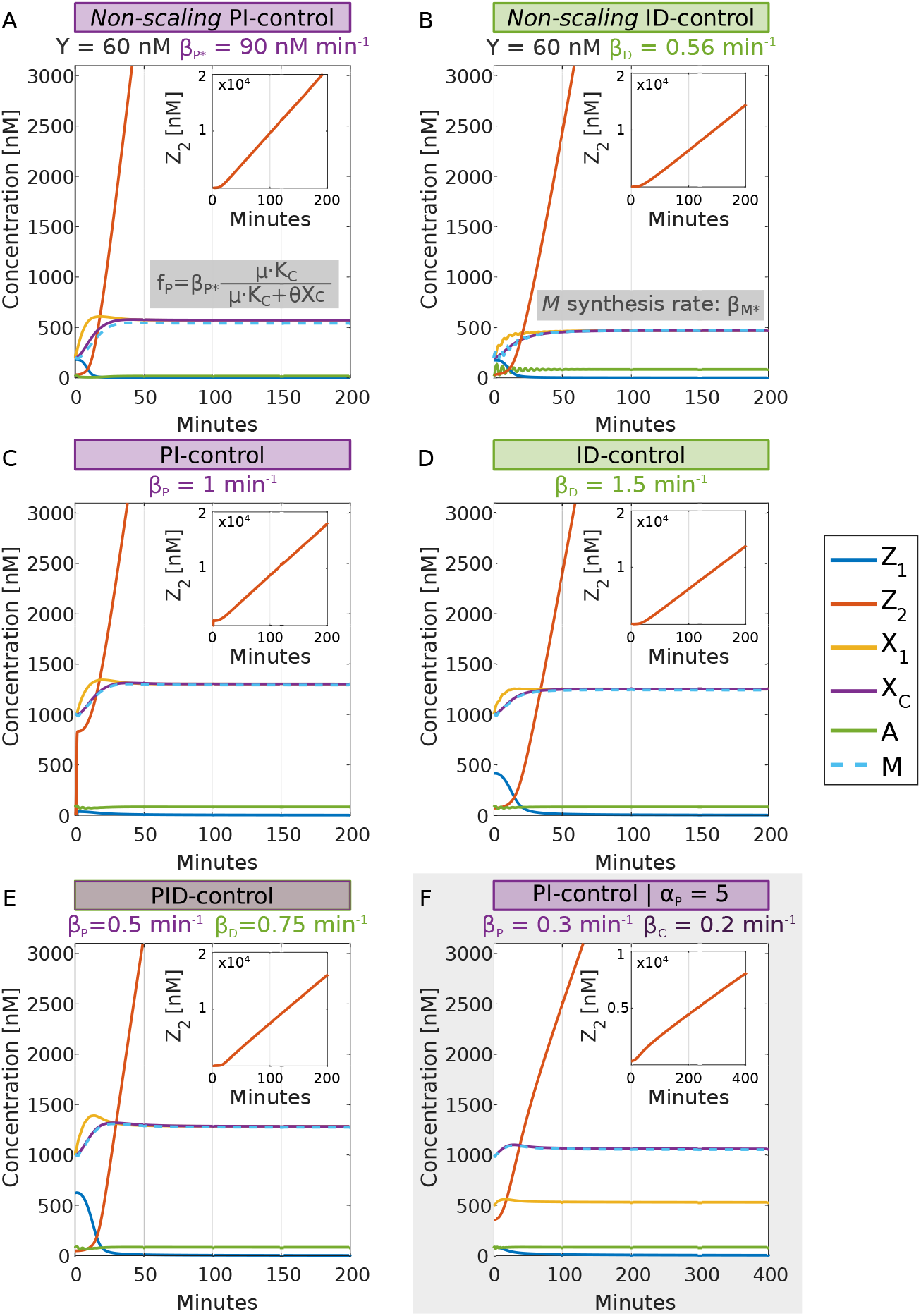
Suboptimal performance occurs when the Proportional or Derivative controllers are too strong. *Z*_2_ levels increase linearly over time when the (A,C) proportional, (B,D) derivative, or (E) a combination of these are too strong and the production of *Z*_1_ is too small to compensate for the production of *Z*_2_. This occurs, for example, for low *Y* set-point values. In all plots the state variables reach a steady-state except for *Z*_2_. (A,B) Plots of all state variables for controllers in which the proportional (A) or derivative (B) terms do not scale with the setpoint *Y*. In these cases, *Y* = 60 nM is too small for the strength of the proportional and derivative functions, respectively (see Fig. 2A and Fig. 3B). (C-E) Plots of all state variables for controllers PI and ID designed in this work, in which controller scales with setpoint *Y*. When *β*_*P*_ and *β*_*D*_, or a combination of these, are too high, loss of integral function is also observed. (F) When *α*_*P*_ ≠ *μ*/*θ* is too high (see Figure S3C), loss of integral control also occurs. See Table S1 for parameter values used in each simulation.

**Figure S3.**
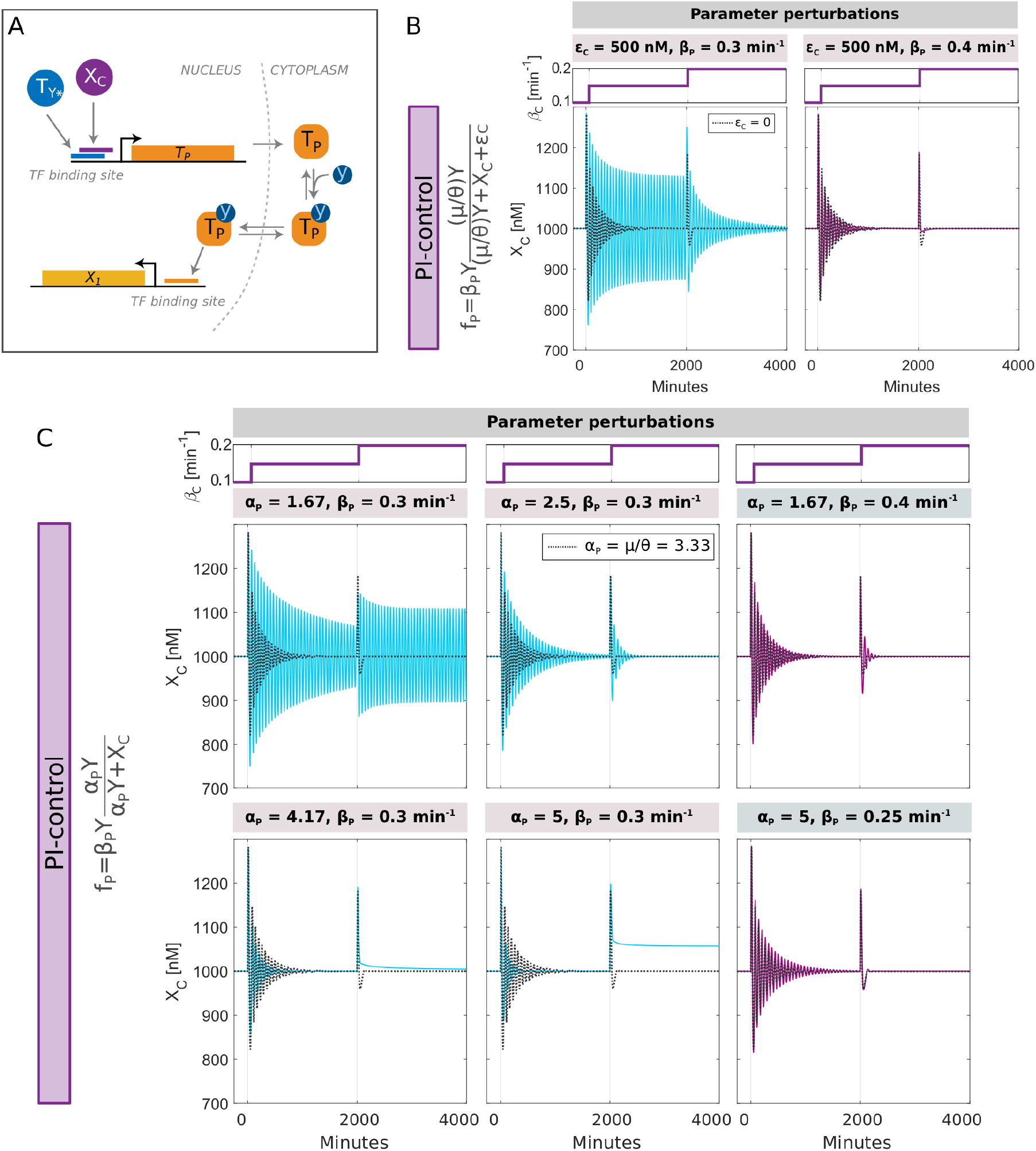
Proportional function realization and non-ideal cases. (A) Proposed experimental realization of the proportional control function 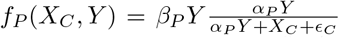 (Eq. s5). (B) [Left panel] Time dynamics of output *X*_*C*_(*t*) following a change in process parameter *β*_*C*_ for *Y* = 300 nM in a model with a PI controller when *α*_*P*_ = *μ*/*θ* = 3.33 and *ϵ*_*C*_ = 500 nM. The example with *ϵ*_*C*_ = 0 is shown on the same plot as a dotted line for comparison. [Right panel] Same plot with *ϵ*_*C*_ = 500 nM, but for an increased value of *β*_*P*_ from *β*_*P*_ = 0.3 min^−1^ to *β*_*P*_ = 0.4 min^−1^. In general, increasing *ϵ*_*C*_ weakens the proportional feedback, but this effect can be compensated for by increasing the value of the proportional weight *β*_*P*_. (C) Time dynamics of output *X*_*C*_(*t*) following a change in process parameter *β*_*C*_ for *Y* = 300 nM in a model with a PI controller when *α*_*P*_ ≠ *μ*/*θ* and *ϵ*_*C*_ = 0 nM. In all cases, the example with *α*_*P*_ = *μ*/*θ* = 3.33 is shown as a dotted line for comparison. Last column showing different values of *β*_*P*_ illustrates that decreasing *α*_*P*_ weakens the proportional feedback and that this effect can be compensated for by increasing the value of *β*_*P*_. However, for large enough *α*_*P*_ (e.g. *α*_*P*_ = 5), the integral controller is compromised (see Figure S2F for *Z*_1_ and *Z*_2_ dynamics). But, this again can be compensated for by decreasing *β*_*P*_. This illustrates the iterative design process that should be undertaken in these systems. See Table S1 for parameter values used in each simulation.

**Figure S4.**
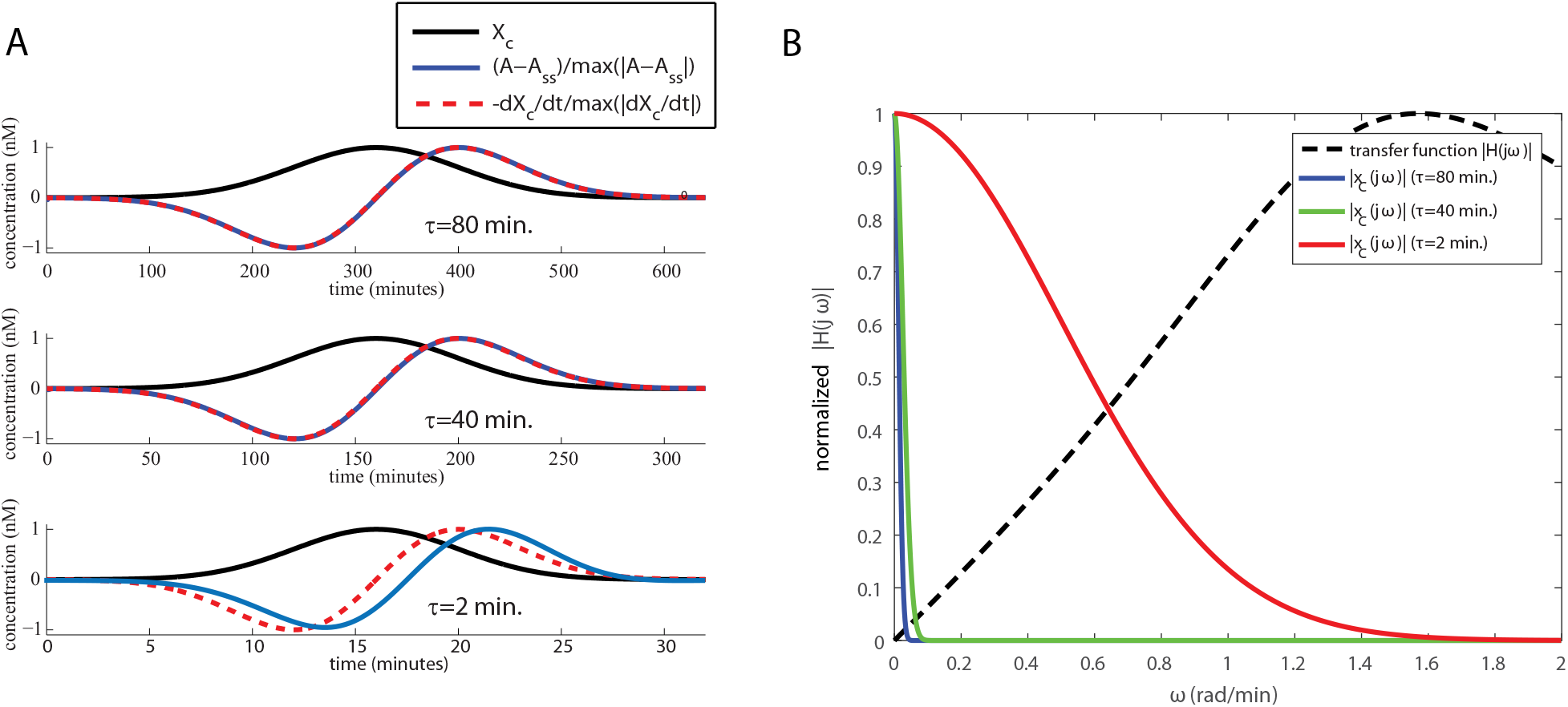
Illustration of design choices and parameter constraints of derivative motif. (A) Simulation of Eqs.(13–14) with parameter values *γ*_*A*_ = 31.4 min^−1^, 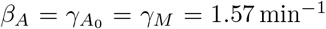, *β*_*M*_*Y* = 1.57 nM min^−1^. *X*_*C*_ is an input to the derivative motif, and is simulated to be a Gaussian of width *τ* and amplitude 1, 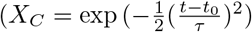, where *τ* = [80, 40, 2] min. Along with *X*_*C*_(*t*) we plot *-dX*_*C*_(*t*)/*dt*/max|*dX*_*C*_(*t*)/*dt*| and (*A*(*t*) – *A*_*ss*_)/max |*A*(*t*) – *A*_*ss*_| where 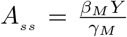, normalized in this fashion for comparison between –*dX*_*C*_(*t*)/*dt* and *A*(*t*). For *τ* = 40 and 80 minutes (i.e. lower frequency content) *A* is accurately representing –*dX*_*C*_/*dt*, but once *τ* is too small (τ = 2 min), *A* starts to deviate from –*dX*_*C*_/*dt*. (B) Frequency domain representation of *X*_*C*_(*s*) and transfer function *H*(*s*) where *s* = *jω* of the derivative motif. For smaller *τ* = 2, *X*_*C*_(*s*) has higher the frequency content that substantially overlap with *H*(*jω*) for the choice of parameters for the derivative motif. This violates the condition that 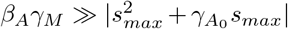 (here *s*_*max*_ ≈ 1.57 rad/min, i.e. the value of *ω* where |*H*(*jω*)| peaks). By contrast, for *τ* = 80 and *τ* = 40, this condition is satisfied.

**Figure S5.**
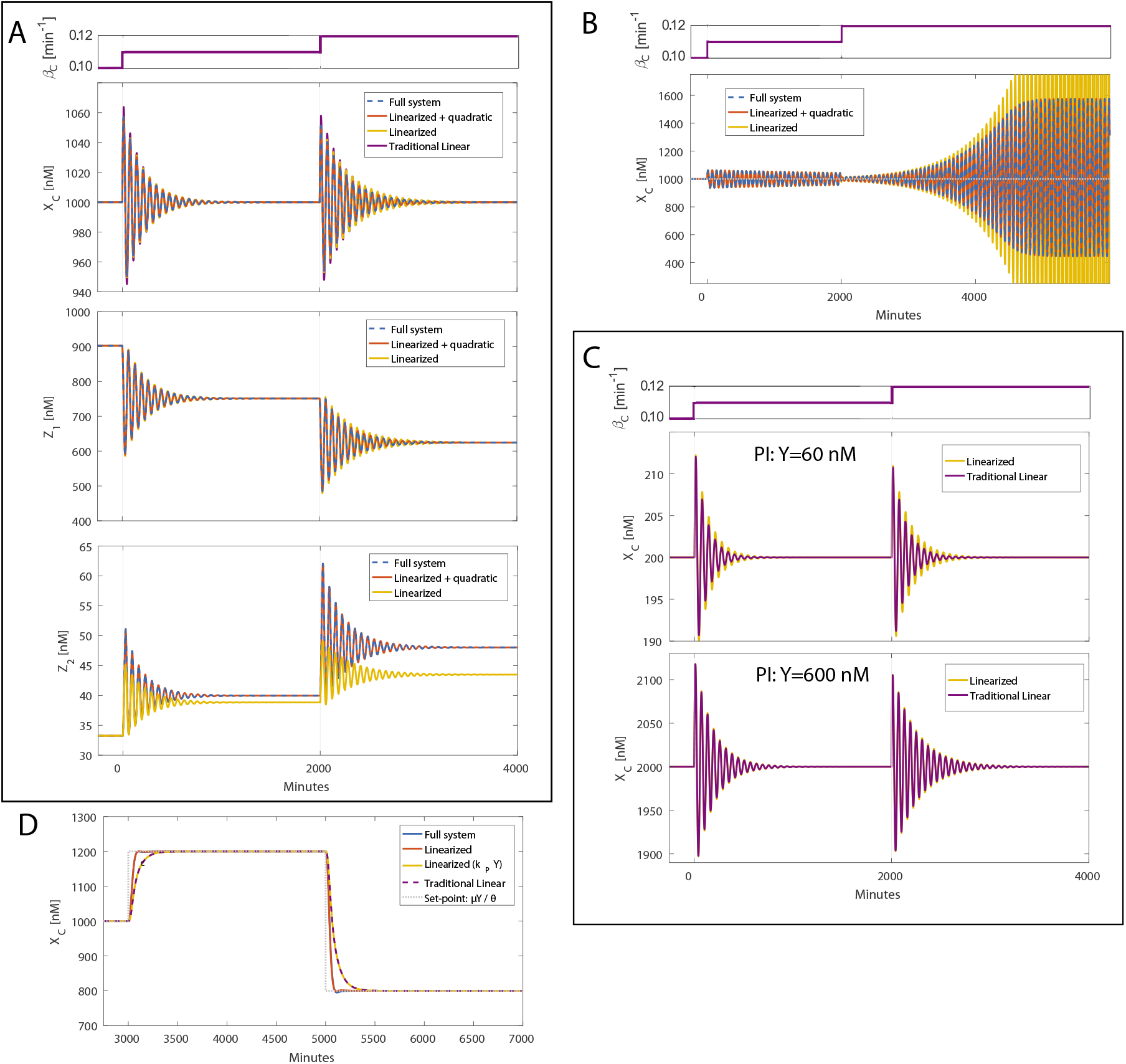
Linearized biochemical PID-control mimics the adaptation dynamics of the nonlinear controller for small to moderate step-changes in the parameter *β*_*C*_ and compares favorably to traditional PID controller under defined parameter regimes. (A) The time dynamics of *X*_*C*_, *Z*_1_ and *Z*_2_ are plotted for the full model (dashed blue, Eqs.(5–8,11–12)), the linearized model (yellow, Eq.(s9)), and the linearized model plus quadratic correction term, *ηz*_1_(*t*)*z*_2_(*t*) (red). See Table S1 for parameter values used in each simulation. The *β*_*C*_ perturbations are shown in the top panels. For *X*_*C*_ dynamics, the traditional PID case (purple) was simulated with just the process (Eq.(5)) and the *X*_1_ equation with exact error terms (Eq.(s15)). (B) Linearized model (yellow) for the integral (I) only case, captures convergent oscillations (first change in *β*_*C*_) and the onset of limit cycle (second change in *β*_*C*_). Adding the quadratic term (red) agrees well with the full solution (dashed blue). (C) *X*_*C*_ dynamics for the linearized biochemical controller and traditional linear controller following step changes in *β*_*C*_ for *Y* = 60 (upper panel) and *Y* = 600 (lower panel). As *Y* increases, the two controllers converge as a result of the convergence of the *k*_*I*_ of the biochemical controller to that of the traditional controller. (D) *X*_*C*_ dynamics for the process with delay (no feedback) from Section 2.4 (Figure 5A). In this case, there is a difference between the traditional PI controller output is not the same as that of the full nonlinear biochemical PI controller and its linearized model. However, changing the proportional terms from 3*k*_*P*_*Y* to *k*_*P*_*Y* makes all the controllers indistinguishable, illustrating a difference between a traditional proportional controller and the biochemical one we design in this work.

